# Discovery of Glycation-Derived Crosslinks at Arginine

**DOI:** 10.1101/2025.07.28.667285

**Authors:** Jeremiah W. Jacob-Dolan, Amy C. Sterling, Morgan E. Brutus, Stefan M. Hansel, Rebecca A. Scheck

## Abstract

Glycation crosslinks account for more than 40% of all known advanced glycation end products (AGEs) and are correlated with many age-related diseases. Despite much interest, crosslinking AGEs (xl-AGEs) remain poorly understood, as they have been challenging to discover, prepare, and quantify. Here we describe a peptide platform that is ideally suited for the study of xl-AGEs, which not only facilitates direct comparisons between the prevalence of known xl-AGEs and other AGEs, but also enables the discovery of previously unknown xl-AGEs. In this study, we use this platform to discover the first known Arg-Arg xl-AGEs, a pair of **<u>m</u>**ethylglyoxal-derived dihydroxy**i**mi**<u>d</u>**azolidine hemi**<u>a</u>**cetal cross**l**ink, or MIDAL, isomers. We show that MIDAL can become the major AGE, exceeding levels of all other AGEs, for substrates in which two Arg glycation sites are optimally positioned. We further demonstrate that MIDAL is readily and reversibly generated in biocompatible conditions, persisting with a half-life of more than three days. We also demonstrate that MIDAL can form in living mammalian cells, suggesting that it has the potential to be a dynamic, physiologically relevant and functional xl-AGE. This work therefore offers important insights about MIDAL formation and describes a versatile platform to enable the study of xl-AGEs under a variety of conditions. We expect that it will be highly useful for further discovery of biologically relevant glycation crosslinks that are yet to be identified.

## Introduction

Covalent protein crosslinks are involved in an impressive range of cellular functions, ranging from strengthening extracellular matrices^1–7^, assisting in bacterial invasion^8^, regulating the cytoskeleton^9^, participating in signaling cascades^10–12^, or stabilizing intramolecular protein folds^13–15^. While some covalent crosslinks are installed or removed by enzymes, others transpire non-enzymatically, often due to oxidative or metabolic stress^16,17^. For example, crosslinking advanced glycation end-products (AGEs) are thought to be a major source of age-related protein damage, especially for long-lived proteins of the extracellular matrix^18,19^. These crosslinking AGEs form spontaneously from endogenous sugars and sugar-derived metabolites, producing a range of harmful impacts including weakened bone strength^20^, increased myocardial stiffness^21^, and the formation of cataracts^22^ or age-related macular degeneration^23^. As the majority of crosslinking AGEs are associated with aging, they are typically thought to accumulate slowly over time^18,19^. However, recent work has shown that some AGE crosslinks form rapidly, even playing a functional role in activation of a bacterial phospholipase^24^ or the antioxidant response^25,26^. Together, these examples suggest that xl-AGEs perform a range of important functions in cellular signaling and human health. However, current efforts to study crosslinking AGEs have so far been unable to facilitate a clear understanding of their prevalence and are poorly suited for discovering new ones^27^.

AGEs are a chemically heterogeneous set of protein post-translational modifications (PTMs) that are formed through the non-enzymatic browning process known as glycation, in which aldehydes or ketones become covalently linked to biomolecules (**Figure 1**)^28–32^. While 43% of reported AGEs are crosslinks, the rest are individual AGEs that modify just a single Arg, Lys, Cys or N-terminus (**Figure 1A**), such as the methylglyoxal-derived hydroimidazolone (MGH-1) isomers^30,33^. Notably, Arg is a major site of glycation, especially by biologically relevant dicarbonyls such as methylglyoxal (MGO) (**Figure 1B**)^34^. Despite the importance of Arg as a glycation site, to date there have been no xl-AGEs reported between two Arg residues. Instead, any known Arg-containing xl-AGEs also include other side chains such as Lys or Cys (**Figure 1C**)^26,28,35,36^. Additionally, compared to non-crosslinking (or mono-) AGEs, relatively little is known about crosslinking (xl-) AGEs. For example, while glucosepane is thought to be a major xl-AGE^32,37^, we have been unable to find any studies that quantify its prevalence relative to common AGEs such as the MGH isomers or CML.

**Figure 1.**
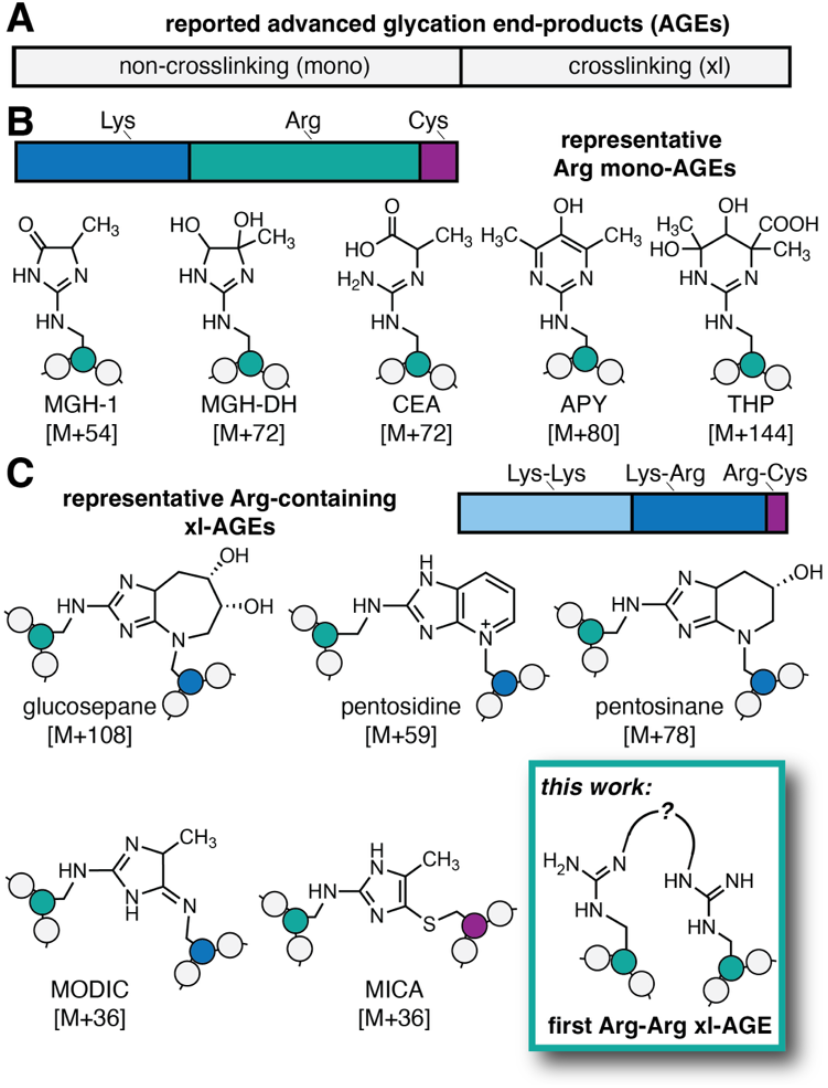
**A)** Of the 40 known advanced glycation end-products (AGEs) that have been reported, almost half (17) are crosslinking AGEs. **B)** For non-crosslinking (referred to as “mono-”) AGEs, the vast majority are found at Lys (9) and Arg (12). Arg is preferentially glycated by the biologically relevant glycating agent methylglyoxal (MGO), forming several AGEs including the methylglyoxal-derived hydroimidazolone isomers (*e*.*g*. MGH-1, shown), dihydroxyimidazolidine (MGH-DH), carboxyethylarginine (CEA), argpyrimidine (APY), and tetrahydropyrimidine (THP). **C)** Known crosslinking (xl-)AGEs are formed mostly between two Lys (9) or a Lys-Arg residue (7), but prior to this work, none have been reported between two Arg. Representative Arg-containing xl-AGEs include glucosepane, pentosidine, pentosinane, methylglyoxal-derived imidazolium crosslink (MODIC) and mercaptomethylimidazole crosslinks between cysteine and arginine (MICA). Here we report the first Arg-Arg MGO-derived crosslink, which we have named MIDAL.

In this work, we set out to develop a peptide-based discovery platform that would enable the study of glycation crosslinks. Here we show that the platform we built is not only suitable for making direct comparisons between known xl-AGEs and mono-AGEs but also enables the discovery of previously unknown xl-AGEs. Specifically, herein we describe the first reported Arg-Arg xl-AGEs, a pair of **<u>m</u>**ethylglyoxal-derived dihydroxy**i**mi**<u>d</u>**azolidine hemi**<u>a</u>**cetal cross**l**ink isomers, which we have named MIDAL. For substrates with optimally positioned Arg, MIDAL becomes the major AGE, surpassing even the formation of non-crosslinking AGEs. We further demonstrate that MIDAL is generated on short time frames and persists for days under mild, biocompatible conditions. Finally, we show that MIDAL can form in living mammalian cells, suggesting that it has the potential to be a major contributor to the glycation landscape. Our findings suggest not only that MIDAL could be a functional xl-AGE, but also there may be additional biologically relevant glycation crosslinks that are yet to be identified.

## Results

Our lab has previously shown that synthetic peptides are useful substrates for evaluating glycation chemistry.^34,38^ In this work, we sought to develop a peptide-based platform that would be particularly well-suited to identify AGE crosslinks, especially those that are Arg-derived. To do so, we envisioned that it would be possible to discern xl-AGEs from any other AGEs by placing a protease recognition site in between glycation sites. Upon glycation, a mixture of mono-AGEs and xl-AGEs would be obtained. However, formation of xl-AGEs would render the sequence recalcitrant to enzymatic cleavage. Therefore, any AGEs remaining on full-length peptides after digestion would indicate that a crosslink had formed (**Figure 2A**). We opted to incorporate Gly and Pro as intervening residues, encouraging conformations in which the two Arg face each other and are primed for crosslink formation. Crucially, this intervening sequence is also a recognition sequence for a Pro-specific endopeptidase^39^. We therefore initiated this study by synthesizing a small library of peptides in which two Arg were separated by a single Pro and a variable number of Gly spacers (**Table S1**). To avoid any potential for glycation at the N-terminus, peptides were acetylated.

**Figure 2.**
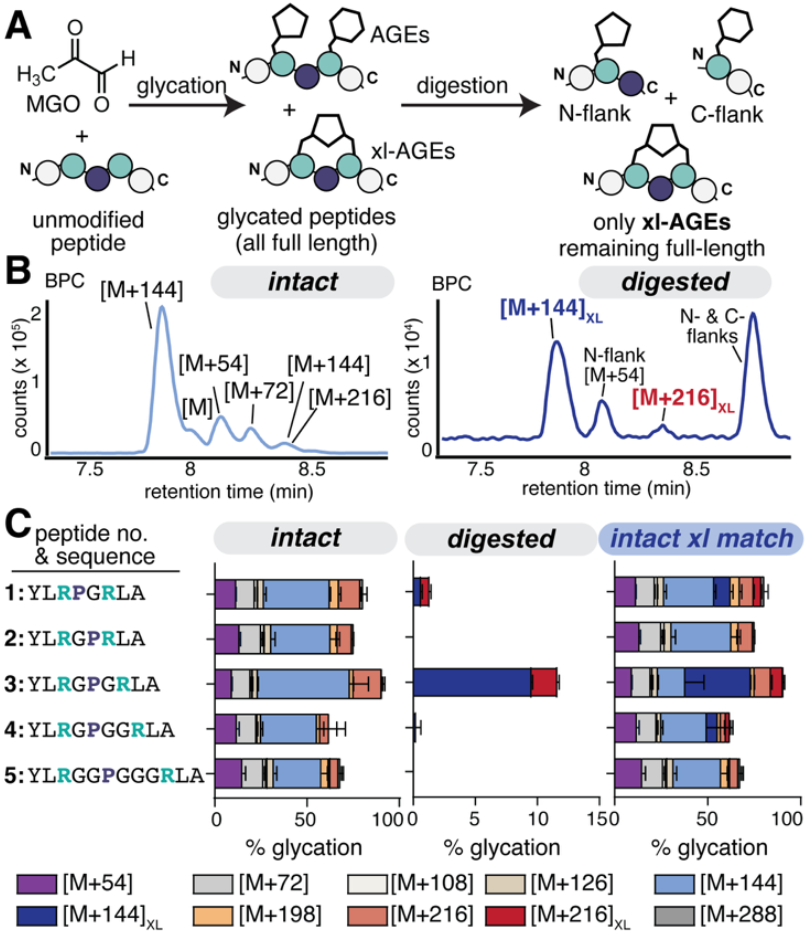
**A)** To evaluate the formation of Arg-Arg glycation crosslinks derived from MGO, we developed a peptide platform in which xl-AGEs could be discerned from all others after cleavage of the peptide backbone by a Pro-specific peptidase. Scheme depicting the use of this platform to screen a small library of peptides (peptides **1** – **5**) in which two Arg (green) were separated by Pro (navy) – Gly linkers of varying lengths. Peptides (1 mM) were treated with 2 mM MGO for 24 h at 37°C in 2X phosphate buffered saline (PBS) at pH 7.4. After quenching the glycation reaction with Tris buffer (intact), a portion of the sample was treated (digested) with the Pro peptidase prior to analysis by liquid chromatography mass spectrometry (LC-MS). **B)** Representative base peak chromatogram (BPC) for MGO-glycated peptide 3 showing intact (*left*) and digested (*right*) samples. **C)** Distribution of AGE adducts observed via LC-MS for peptides **1**–**5** on intact samples (*left*) and on remaining full-length peptides after digestion (*middle*). By matching retention times, intact data was reanalyzed to differentiate xl-AGEs from other AGEs (*right*). Stacked bar graphs are shown with mean ± standard deviation for each adduct. Data is from independent experiments, with n= 9 for peptide 3 and n=3 for all other peptides. Legend: purple [M+54], light gray [M+72], cream [M+108], tan [M+126], light blue [M+144], dark blue [M+144]_XL_, orange [M+198], light red [M+216], dark red [M+216]_XL_, dark gray [M+288].

We used AlphaFold to model the most likely structures for each of our peptides **1**–**5** (**Figure S1**)^40^. In all cases, AlphaFold placed the guanidino groups facing towards each other in the majority of the predicted conformations, with the average inter-guanidino carbon distances ranging from 7-11.5 Å. We also optimized proteolysis conditions and found that the Pro peptidase exhibited highly efficient cleavage on these short peptide substrates, as compared to tobacco etch virus (TEV) on a related set of peptides (**Figure S2**). We confirmed that unmodified peptide was completely cleaved by the Pro peptidase after only 15 min at 30 **°**C. However, we note that the Pro peptidase exhibited modest off-target cleavage for sequences in which an aromatic residue was immediately C-terminal to Arg (**Figure S3**). This behavior guided our design, as we avoided placing Tyr immediately before or after either of the Arg sites.

Next, we evaluated the use of this platform to reveal xl-AGEs by treating each peptide (**1**– **5**, 1 mM) with 2 mM MGO for 24 h at 37 **°**C (**Figure 2A-C**). These conditions are similar to those we have used to assess peptide glycation in the past^38^, though the MGO concentration was doubled to account for the additional Arg residue. After quenching the glycation reaction in Tris buffer, a portion of the sample was further treated with the Pro peptidase prior to analysis by liquid chromatography-mass spectrometry (LC-MS). Using this approach, we observed the expected Arg AGE adducts, including MGH-1 ([M+54]), CEA and MGH-DH ([M+72]), and multiple [M+144] adducts, which likely include THP (**Figure 2B**, *left*.) As there were two potential glycation sites in our substrate, we also observed expected mass changes that aligned with combinations of these adducts, (*e*.*g*. double MGH formation, [M+108]), up to [M+288]. These were observed in roughly similar distributions for all peptides tested (**Figure 2C**, *left*).

By contrast, after digestion, no unmodified peptide remained, and many of these high molecular weight AGEs were no longer present (**Table S2**). Instead, we observed the unmodified N- and C-terminal flanks (*m/z* = 324.1800, *z*=2 (observed); 324.1739, *z*=2 (expected) and 208.5792, *z*=2 (observed); 208.6344, *z*=2 (expected), respectively), liberated after digestion, as well as glycated versions of these fragments (**Figure 2B**, *right*). Of particular interest, however, were two species (*m/z* = 594.8244 and 630.8343, respectively (*z*=2)), both of which were greater than the mass of the unmodified parent peptide we began with (*m/z* = 522.8034, *z*=2). These results suggested that our platform is successfully able to differentiate xl-AGEs from others that form (**Figure 2C**, *middle*).

Although we had predicted that only xl-AGEs would be remaining as full-length (or greater) masses after digestion, we also envisioned two possibilities for this behavior. One possibility was that xl-AGEs would be inert to the Pro peptidase; in this case, we expected to see identical retention times in both intact and digested samples. The other possibility was that the backbone would still be clipped during digestion, leaving a full-length xl-AGE peptide with the addition of water. Careful inspection of retention times (r.t.) and mass changes observed in both intact and digested samples revealed that the observed xl-AGEs were identical in both treatments. Specifically, one xl-AGE adduct ([M+144]_XL_) eluted at r.t. = 7.855 ± 0.012 min with *m/z* =594.8244, *z*=2 in both intact and digested samples. The other xl-AGE ([M+216]_XL_) eluted as a mixture of isomers with retention times at 8.180 ± 0.009 and 8.381 ± 0.004 min, both with *m/z* = 630.8343, *z*=2 that were identical in digested samples. These results suggest that xl-AGEs are indeed resistant to Pro peptidase cleavage, consistent with past studies evaluating proteolysis of cyclic or other constrained peptides.^41^

Having determined the exact retention time for each of the two xl-AGEs in the intact samples, it was possible to perform superior quantification that enabled us to directly compare the relative levels of xl-AGEs to mono-AGEs (**Figure 2C**, *right*). We found that only peptide **3** (Ac-YLRGPGRLA) led to substantial levels of xl-AGE formation, with 40.2 ± 4.8% of [M+144]_XL_ and 7.1 ± 0.9% of [M+216]_XL_. Peptide **3** generated fivefold the levels of xl-AGE formation compared to peptide **1** (8.8 ± 1.4%), and almost ten times the levels observed for peptide **4** (5.8 ± 8.2%). Additionally, no xl-AGEs were observed for peptides **2** or **5**, suggesting that the distance between the two Arg residues is likely to play a major role in the amount of crosslink formed (**Figure 2C, Figure S1**). Moreover, the [M+144]_XL_ was the highest abundance adduct out of all AGEs observed for peptide **3**.

To ensure that the [M+144]_XL_ crosslink was not an artifact of the Pro peptidase, we also synthesized a peptide **3** variant that could be digested using a photolabile linker rather than enzymatic cleavage (**Figure S4**). After photocleavage, in addition to the single AGEs formed on the photocleaved fragments, we observed that only two masses remained at or above that of the parent peptide, which matched with the [M+144]_XL_ and [M+216]_XL_ adducts that we found using the Pro peptidase protocol. Taken together, these experiments confirm that the [M+144]_XL_ adduct is a previously unknown MGO-derived crosslinks that form between two Arg. We further suspect that the [M+216]_XL_ is an additional MGO addition (either MGH-DH or CEA) on peptides already containing [M+144]_XL_, as has been previously suggested for related xl-AGEs^42^.

While there have been no other reported Arg-Arg AGE crosslinks, and no xl-AGEs reported with mass changes of [M+144] or [M+216], there are other known MGO-derived crosslinks that include Arg at one of the reactive sites. To further confirm that we were observing authentic crosslink formation, we therefore sought to use our peptide platform to confirm the presence of known crosslinks. In particular, we focused on two known MGO-derived crosslinks: an Arg – Cys crosslink (MICA), and an Arg – Lys crosslink (MODIC) (**Figure 3A**). To do so, we prepared peptide substrates that resembled peptide **3**, but included a single point mutation changing the Arg at position 3 to a cysteine (peptide **6**) or a lysine (peptide **7**).

**Figure 3.**
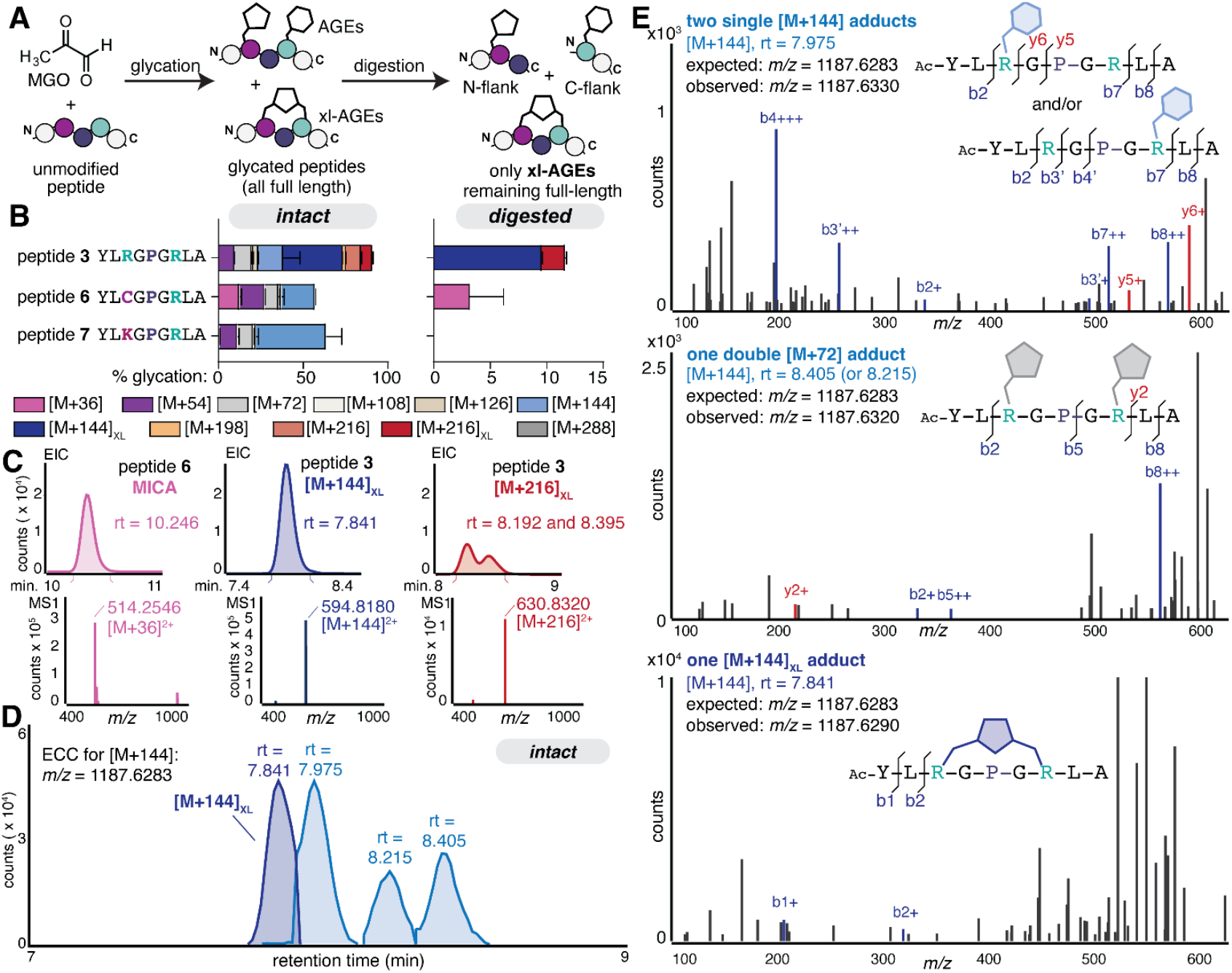
**A)** To confirm crosslink formation, variations of peptide **3** were prepared with Cys (peptide **6**) or Lys (peptide **7**), which are known to form the crosslinks MICA or MODIC, respectively, when treated with MGO. Scheme depicting the treatment of peptides (1 mM) with 2 mM MGO for 24 h at 37**°**C in 2X phosphate buffered saline (PBS) at pH 7.4. After quenching the glycation reaction with Tris buffer (intact), a portion of the sample was treated (digested) with the Pro peptidase prior to analysis by LC-MS. **B)** Distributions of AGE adducts observed via LC-MS on peptides **3, 5** and **6** for intact samples (*left*) and on remaining full-length peptides after digestion (*right*). By matching retention times, intact data was reanalyzed to differentiate xl-AGEs, as shown in the intact AGE distributions (*left*). Stacked bar graphs are shown as the mean ± standard deviation for each adduct. Data is from independent experiments, with n= 9 for peptide **3** and n=3 for all other peptides. Legend: pink [M+36], purple [M+54], light gray [M+72], cream [M+108], tan [M+126], light blue [M+144], dark blue [M+144]_XL_, orange [M+198], light red [M+216], dark red [M+216]_XL_, dark gray [M+288]. **C)** Representative extracted ion chromatograms (EIC) for crosslinking adducts and corresponding mass spectra. **D)** Representative extracted compound chromatograms for four discrete [M+144] isomers from the same intact sample of peptide **3** treated with MGO. **E)** MS^2^ spectra for [M+144] isomers (precursor ion *m/z* = 1187.6283). Diagnostic b and y ions were found for the isomer at observed at retention times of 7.975 min (b2, b3’, b4’, y6, y5, b7 and b8), suggesting a single THP (or other [M+144] mono-AGE) (*top*), and for those at 8.405 min and 8.215 min (b2, b5 and y2) suggesting two [M+72] mono-AGEs, one on each Arg (*middle*). For the [M+144]_XL_ isomer observed at a retention time of 7.841 min, no diagnostic b and y ions were identified, providing further evidence of crosslink formation (*bottom*).

Unlike the xl-AGEs we identified for Arg-only sequences, both MICA and MODIC crosslinks have a unique mass change ([M+36]) that is conveniently tracked even without digestion. When treated with MGO, we found that both peptides **6** and **7** were highly AGE modified (57.5 ± 2.3% and 63.4 ± 9.9% respectively), though peptide **3** produced the highest level of overall glycation (90.6 ± 8.1%). Prior to digestion, we observed an [M+36] mass adduct for peptide **6** (12.6 ± 1.0%) suggesting that MICA, is able to form under the conditions used in this study. For peptide **7**, an [M+36] adduct was observed in vanishing quantities (0.1 ± 0.2%) but only in intact samples, suggesting minimal MODIC formation that was not sufficient to survive digestion (**Figure 3B**). This difference may be attributed to the fact that MGO reacts quickly with both Arg and Cys, but reacts far more slowly with the Lys ε-amine.^43^ This difference could also be due to a conformational effect, as the average AlphaFold predicted distances between the terminal side chains for peptides **3** and **6** were quite similar (9.49 ± 3.35 Å vs 9.2 ± 2.18 Å, respectively), while the distance for peptide **7** was far greater (13.74 ± 3.36 Å) (**Figure S1**).

Representative extracted ion chromatograms (EIC) for all crosslinks observed show that MICA and the [M+144]_XL_ elute as single, well-resolved peaks, but the [M+216]_XL_ appears as multiple, broad peaks and in far lower quantities (7.1 ± 0.9%) of total peptide volume compared to [M+144]_XL_ (40.2 ± 4.8%) (**Figure 3C**). For these reasons, we focused our attention on the [M+144]_XL_ and its comparison to MICA. Specifically, the discrete mass change for MICA allowed us to directly compare levels of [M+36] in both intact and digested samples. We found that MICA (r.t. = 10.260 ± 0.009 min; m/z = 514.2574, z=2) was the only AGE to survive digestion for peptide **6** (**Figure 3B**). After digestion, there was a substantial loss of absolute counts, which were decreased by an order of magnitude (**Figure S5**). This matches the behavior we observed for [M+144]_XL_ on peptide **3**. We suspect that the loss of counts after digestion could be due to either off-target cleavage from the peptidase, or on-target cleavage that destabilizes the crosslink. Nonetheless, these data support the use of the Pro peptidase to confirm the exact mass and retention time of each xl-AGE, with any quantification relative to other AGEs taking place on the intact (undigested) sample. By doing so, we found that the [M+144]_XL_ we identified appears to be of greater prevalence than MICA for comparable substrates (40.2 ± 4.8% vs 12.6 ±1.0%, respectively) (**Figure 3B, C**). Unless otherwise noted, all subsequent AGE distributions reported are based on quantification from intact samples, with retention time matching between intact and digested samples performed in a pairwise manner.

Unlike MICA and MODIC, the new crosslink that we uncovered does not have a unique mass. An [M+144] AGE is degenerate with multiple possibilities, including the formation of tetrahydropyrimidine (THP), other AGEs with a mass change of [M+144], or the formation of [M+72] (MGH-DH or CEA) on both Arg sites. Accordingly, we were able to observe four [M+144] adducts with discrete retention times during our intact experiment (**Table S2, Figure 3D**). As further confirmation of crosslink formation, we performed targeted tandem mass spectrometry (MS^2^) analysis on [M+144] species (precursor ion *m/z =* 594.8, isolation width = 4 *m/z*). For the peak eluting at 7.913 ± 0.104 min (observed in a representative sample at 7.975 min), the resulting MS^2^ spectra were consistent with the formation of a single THP (or other [M+144] mono-AGE) on either Arg, with both species co-eluting. Diagnostic ions included b2 and y6 ions, showing modification of the first Arg (at position 3), and the b4’ and b7’ ions showing modification of the second one (at position 7) (**Figure 3E**, *top*). At a retention time of 8.412 ± 0.009 (**Figure 3E**, *middle*) and 8.231 ± 0.014 min (observed at 8.405 min and 8.215 min, respectively), the b2, b5, and y2 ions are diagnostic for another [M+144] species that has two [M+72] modifications, one at each Arg. Finally, in the MS^2^ analysis for the [M+144]_XL_ at 7.855 ± 0.012 min (**Figure 3E**, *bottom*) (observed at 7.841 min), no diagnostic ions were observed. The only b or y ions that could be assigned were for b1 or b2, which fall outside of the crosslink itself. We expect that this occurs because MIDAL forms a cyclic peptide that prevents the expected fragmentation that would usually result from collision-induced fragmentation at amide bonds during conventional MS^2^. While some known intermolecular crosslinks between linear peptides^44,45^ or C- to N-cyclized peptides can be sequenced by MS^2^ with non-conventional fingerprints,^46,47^ we have not found any examples of MS^2^ analysis in which a non-amide crosslink is responsible for cyclization. Thus, we interpret the inability to identify any diagnostic ions as further evidence of crosslink formation.

Although many AGE crosslinks have been reported, none have a mass change of [M+144], suggesting that the one we discovered has a novel structure. To aid in determining the [M+144]_XL_ structure, we used two MGO derivatives (**Figure 4A**). First, we used a deuterated version of MGO (D4-MGO), in which all four protons are replaced with deuterium. Experiments using deuterated MGO exhibited no exchange of protons during crosslink formation, generating an [M+152]_XL_ adduct that differed by the expected 8.00 Da from the [M+144]_XL_. We also used glyoxal (GO) to show that a methyl group was not required to form the crosslink, as we observed an [M+116]_XL_ adduct that corresponds to the [M+144]_XL_ obtained for MGO (**Figure 4B,C**).

**Figure 4.**
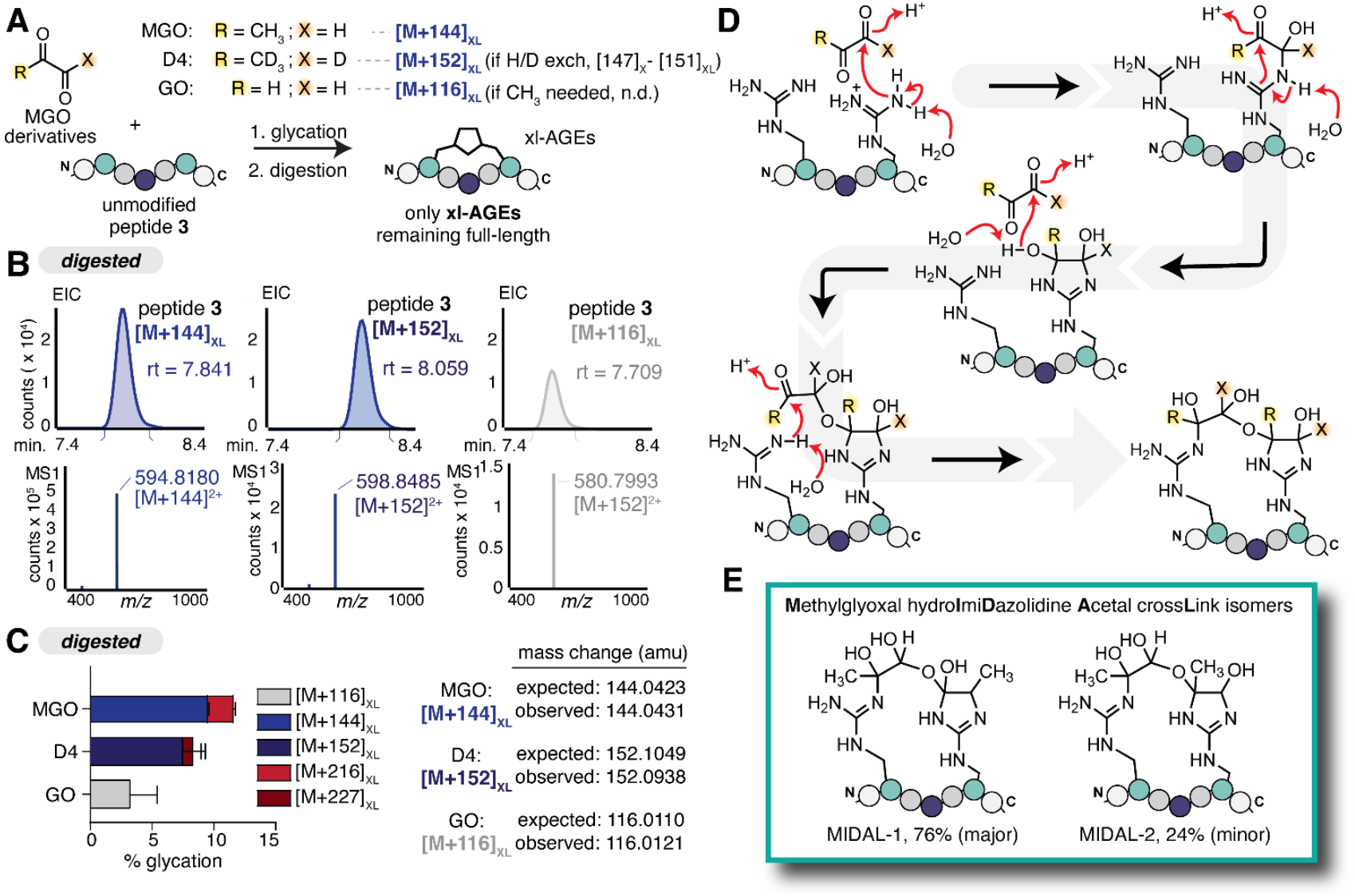
**A)** Glycation reactions were performed by incubating 1 mM peptide **3** with either 2 mM MGO, 2 mM deuterated (D4)-MGO, or 5 mM glyoxal (GO) in 2X PBS at pH 7.4 for 24 h at 37 °C, followed by digestion with the Pro peptidase to track formation of xl-AGEs. **B)** Representative extracted ion chromatograms (EIC) and corresponding MS spectra for xl-AGEs remaining at or above the parent mass after glycation and subsequent digestion. **C)** AGE distributions for digested samples after glycation. Expected and observed mass changes for each MGO derivative are shown. Stacked bar graphs show the mean ± standard deviation for each xl-AGE adduct, quantified on the digested samples. Data is from independent experiments, with n= 9 for peptide **3** treated with MGO and n=3 for D4-MGO and GO treatments. Legend: light gray [M+116]_XL_, blue [M+144]_XL_, dark blue [M+152]_XL_, red [M+216]_XL_, dark red [M+227]_XL_. **D)** Proposed mechanism for [M+144]_XL_ formation that is consistent with observations in panels B & C. **E)** NMR characterization supports the assignment of **<u>m</u>**ethylglyoxal-derived dihydroxy**i**mi**<u>d</u>**azolidine hemi**<u>a</u>**cetal cross**l**ink (MIDAL) isomers (MIDAL-1, 76%; MIDAL-2, 24%). Structural characterization can be found in Supplementary Figures S6 and S7.

While many reported AGEs suggest involvement of the methyl group and/or proton exchange, prior work outside of the glycation literature has suggested that dihydroxyimidizolidines can condense with an additional carbonyl, forming an acetal^48,49^. Though past work has focused on this kind of linkage as a stable, standalone modification, we envisioned that similar chemistry could be at play in generating a crosslink. However, acetal formation between two MGO [M+126] would be inconsistent with the observed mass change. As a result, we instead considered structures and mechanisms in which no waters were lost. This led us to consider an alternative pathway where the nearby Arg catches and thereby stabilizes the hemiacetal intermediate. Such a mechanism would be consistent with our findings when using the MGO derivatives, as all MGO protons and the methyl group do not participate in crosslink formation (**Figure 4D**). Finally, using a combination of 1D and 2D NMR, we were able to confirm the structure of the [M+144]_XL_ crosslink, which involves a **<u>m</u>**ethylglyoxal-derived dihydroxy**i**mi**<u>d</u>**azolidine hemi**<u>a</u>**cetal cross**l**ink between Arg, which we have named MIDAL (**Figure 4E, Figures S6 & S7**).

To provide insight into the potential for MIDAL to form in cells, we evaluated the conditions that promoted its formation. We subjected peptide **3** to a variety of different glycation reaction conditions, scanning reaction time, temperature, MGO concentration, and pH (**Figure S8**). These data showed that MIDAL formation was maximal at neutral pH (7.4) and 37 **°**C. Further, our results suggest that MIDAL is likely to form quickly, as it reaches its maximal levels in our *in vitro* system around 24 h (40.2 ± 4.8%), before it begins rearrangement into mono-AGEs such as MGH-DH and CEA. After 24 h of additional MGO incubation, MIDAL levels drop to 20.3 ± 3.1%. To determine the stability of MIDAL in the absence of MGO, we incubated purified MIDAL-modified peptide 3 at 37 °C in PBS at pH 7.4. Under these conditions, 43.7 ± 3.0% of MIDAL survived after a 4-day incubation (**Figure S9**). The removal of MGO from further incubation showed a modest increase in the stability of MIDAL, revealing its potential to be modulated by the available level of glycating agent (in this case MGO). Using these conditions we calculated the half-life of MIDAL to be 3.3 days which is longer than early AGEs such as MGH-DH, (t_1/2_ = 1.8 days)^50^ but shorter than AGEs such as fructosyl-lysine (t_1/2_ = 25 days) or the MGH isomers (t_1/2_ = 12 days)^51,52^. Collectively, these data demonstrate that MIDAL fits the profile of a dynamic and biologically relevant AGE, which not only forms readily, but also persists for several days at physiological pH and temperatures.

Having both confirmed the structure of MIDAL and established that it forms under mild conditions, we sought to determine if it had the potential to form in a cellular environment. To do so, we transiently transfected HEK293T cells with a plasmid in which the peptide **3** (-YLRGPGRLA, GFP-**3**), peptide **6** (-YLCGPGRLA, GFP-**6**) or peptide **7** sequence (-YLKGPGRLA, GFP**-7**) was fused at the C-terminus of green fluorescent protein (GFP) (**Figure 5A**). In addition to the internal Pro peptidase recognition site, these C-terminal sequences were connected to GFP through a linker sequence containing a tobacco etch virus (TEV) protease cleavage site (**Figure 5B**). We confirmed that all three variants were comparably expressed in HEK-293T cells after 24 h of transient transfection (**Figure 5C**). Next, to observe crosslink formation, we incubated transfected cells with or without 10 mM MGO for 2 h (*Treatment A*, **Figure 5A**). These treatment conditions were used to maximize crosslink formation while also minimizing MGO toxicity. Specifically, we found that even after 2 h of incubation with these high concentrations of MGO, >80% of cells remained viable, consistent with previous findings^53^.

**Figure 5.**
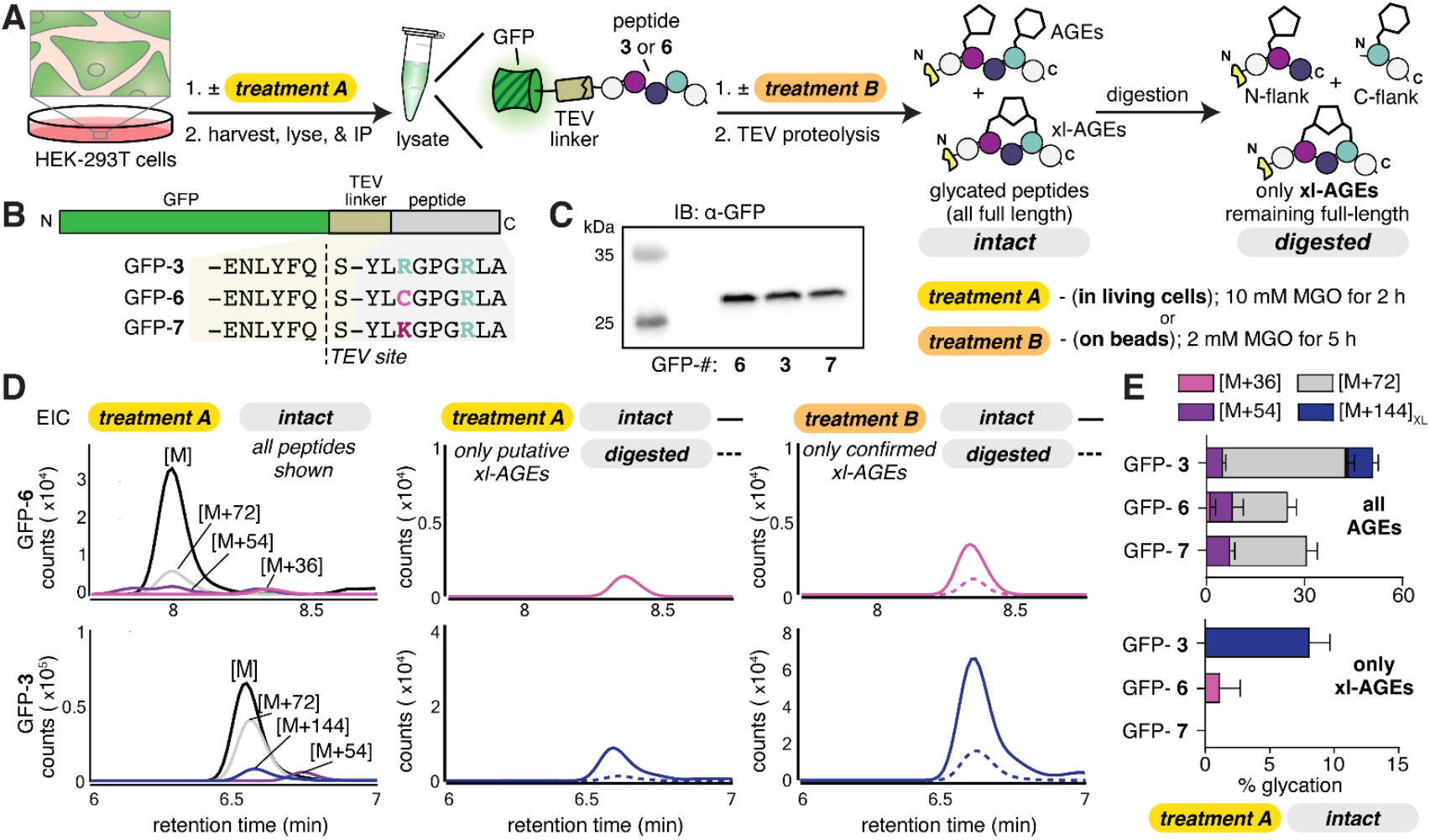
**A,B)** Scheme depicting the workflow used to assess MIDAL formation in cells. Briefly, HEK-293T cells were transfected with plasmids encoding the green fluorescent protein (GFP) fused to peptide **3, 6**, or **7**, linked via the tobacco etch virus (TEV) protease recognition site (GFP**-3, -6**, or **-7**, respectively). *Treatment A:* cells were treated with or without 10 mM MGO for 2 h prior to lysis. *Treatment B:* after lysis and immunoprecipitation, GFP-bound beads were treated with 2 mM MGO for 5 h. After either MGO treatment protocol, peptides were released from resin using TEV proteolysis. The resulting peptides were analyzed by LC-MS (intact) or subsequently digested with the Pro peptidase (digested). **C)** Western blot analysis, probing with α-GFP antibodies, revealed that each GFP variant was expressed comparably in in HEK-293T cells. **D)** Representative EICs for peptides released from GFP-**6** (top row) or GFP**-3** (bottom row) after MGO treatment. *Left*, overlay of all peptide species observed in intact samples from treatment A. *Middle*, overlay of putative crosslinked species in intact (solid) and digested (dashed) samples from treatment A. *Right*, overlay of confirmed crosslinked species in intact (solid) and digested (dashed) samples from treatment B. **E)** AGE distributions for intact samples obtained from MGO treatment A. Stacked bar graphs showing all AGEs (top) or only xl-AGEs (bottom) are shown as the mean ± standard deviation for each adduct. Data is from independent experiments, with n= 3. Legend: pink [M+36], purple [M+54], gray [M+72], blue [M+144]_XL_.

Following MGO treatment, cells were harvested, and GFP-peptide fusions were immunoprecipitated. Taking advantage of the TEV cleavage site, we then performed an on-bead TEV digestion to release the peptides of interest, which were analyzed by LC-MS. We found that GFP-**3** led to a mixture of AGEs (50.5 ± 4.3% modification), including [M+54], [M+72], [M+126], and a single peak with an [M+144] adduct (15.8 ± 3.2% modification) (**Figure 5D**,**E**). Furthermore, this [M+144] adduct was the only one remaining after treatment with the Pro peptidase. Notably, GFP-**6** also exhibited [M+54] and [M+72] AGEs, lacked [M+126] and [M+144], and instead had the [M+36] AGE, corresponding to MICA. However, when using this crosslink as a positive control, we found that we were unable to observe it after treatment with the Pro peptidase, likely due to the low levels (1.1 ± 1.5%) of MICA prior to digestion. We also noted that, consistent with our results *in vitro*, no crosslink formation was observed using GFP-**7**, though we observed [M+54] and [M+72] mono-AGEs.

To ensure that the [M+144] remaining after digestion was indeed MIDAL, we used an alternative MGO treatment protocol (*Treatment B*, **Figure 5A**) that would allow us to both maximize xl-AGE levels and match all crosslink retention times for MGO-treated cells, including our positive control. Specifically, in the alternative protocol, we performed on-bead MGO treatment (2 mM, 5 h), after GFP immunoprecipitation, but prior to elution with TEV protease. After this treatment, we used TEV protease to liberate the C-terminal peptides. This MGO treatment protocol led to substantially increased levels of glycation, with no change in the identities of the AGEs observed. For GFP-**3**, overall glycation levels jumped from 50.5 ± 4.3% with treatment A to 88.9 ± 4.5% for treatment B. Again, only a single peak with an [M+144] adduct was observed, but at a far greater level accounting for more than half of total glycation (56.4 ± 4.9% of total glycation). For GFP-**6**, glycation levels increased from 24.6 ± 2.9% with treatment A to 66.1 ± 1.1% for treatment B. This increase in glycation enabled us to confirm the retention time for any xl-AGEs that remained following Pro peptidase digestion. After digestion, we confirmed that the [M+36] adduct persisted, with identical retention times (**Figure 5D, Table S2**). This same approach was used to demonstrate that the [M+144] adduct we observed for the peptide released from GFP-**3** also persisted even after digestion, supporting its assignment as MIDAL (**Figure 5D, E**). Taken together, these results provide clear evidence that MIDAL not only forms readily in biological systems, but also that it may be prevalent compared to many known AGEs, not only to other AGE crosslinks.

## Conclusion

Despite some recent progress^53,54^, it remains extraordinarily challenging to monitor xl-AGE formation. Accordingly, most prior efforts to discover xl-AGEs have primarily relied on isolation (and subsequent characterization) from aged protein or tissue samples^29,33,35^. While some xl-AGEs have been observed serendipitously^24,25^, most recent efforts to facilitate their study have focused on the chemical synthesis of these known crosslinks to generate affinity reagents^36,55,56^. However, none of these approaches enable direct comparisons between levels of xl-AGEs and other AGEs and are poorly suited for discovering new xl-AGEs. To address this need, here we have described a peptide-based platform that enables the study of glycation crosslinks. This platform allowed us not only to directly compare the prevalence of xl-AGEs to other AGEs, but also to discover the first known arginine-arginine crosslink formed by methylglyoxal, which we call MIDAL. MIDAL contains a novel crosslink structure involving a surprisingly stable hemiaminal and hemiacetal. Our results show that MIDAL formation likely depends on the inter-Arg distance, as varying this distance affects the ratio of crosslinking observed relative to other AGEs. We further demonstrate that MIDAL forms readily and, though its formation appears to be reversible, it also persists for days under mild, biocompatible conditions. We therefore expect that it could be an important, but so far overlooked, AGE.

Our work also shows that MIDAL can indeed form in cellular systems, further suggesting it may play a biological role that aligns more closely with crosslinks that participate in functional signaling pathways^24,26^ rather than those that are considered markers of long-term damage^20–22^. Like any PTMs, xl-AGEs may alter local chemical properties and/or recruit new binding partners. However, MIDAL is also likely to impose new constraints on protein structure, dynamics, and interactions. Furthermore, our findings imply that the extent of glycation-derived crosslinking may be underestimated. For substrates with optimally positioned Arg, which we crudely estimate to be around 9.5 Å, MIDAL becomes the major AGE, surpassing even the formation of non-crosslinking AGEs. Our future work will therefore focus on better defining the optimal inter-Arg distance to streamline the identification of authentic MIDAL substrates.

Due to the degenerate mass of MIDAL with other reported AGEs formed from MGO, including THP and double CEA (or MGH-DH) modifications, at present it is intractable to profile MIDAL using unbiased proteomics workflows, which require knowledge about expected mass changes and sites for dynamic modifications. We suspect that other crosslinks may have similarly eluded previous detection, being misattributed as another adduct with the same mass change. While we plan to develop tools and reagents that enable the detection of MIDAL substrates using proteomics, we also expect to use our peptide platform to evaluate the formation of other xl-AGEs, and their relative abundances compared to more commonly studied AGEs such as CML or MGH isomers. These studies will provide critical insights into the chemistry of glycation crosslinking events and uncover their resulting biological consequences.

## ASSOCIATED CONTENT

### Supporting Information

This material is available free of charge via the Internet at http://pubs.acs.org.”

## AUTHOR INFORMATION

### Author Contributions

J.W.J.-D. designed the project, performed all experiments and analysis, and prepared the figures and manuscript. M.E.B assisted with NMR acquisition and assignment of MIDAL structure. A.C.S synthesized deuterated MGO using protocols developed by S.M.H. R.A.S designed the project, directed the experimental design and analysis, and prepared the figures and manuscript.

### Notes

The authors declare no competing interests.

## ABBREVIATIONS

AGE: advanced glycation end product
PTM: post-translational modifications
MGH-1: methylglyoxal-derived hydroimidazolone
MGO: methylglyoxal
MGH-DH: dihydroxyimidazolidine
CEA: carboxyethylarginine
APY: argpyrimidine
THP: tetrahydropyrimidine
Xl: crosslinking
MODIC: methylglyoxal-derived imidazolium crosslink
MICA: mercaptomethylimidazole crosslinks between cysteine and arginine
MIDAL: methylglyoxal-derived dihydroxyimidazolidine hemiacetal crosslink
TEV: tobacco etch virus
LC-MS: liquid chromatography mass spectrometry
BPC: Base peak chromatogram
R.T: retention time
PBS: phosphate buffered saline
EIC: extracted ion chromatogram
GO: glyoxal
D4-MGO: deuterated methylglyoxal
GFP: green fluorescent protein
IP: immunoprecipitation

