## Supporting Information for "Discovery of Glycation-Derived Crosslinks at Arginine"

**Materials and Methods** 2

**Supporting Figures and Tables**
**Table S1.** Peptide masses and retention times 8
**Table S2.** AGE adducts observed 9
**Figure S1.** AlphaFold predictions of peptides 1-7 10
**Figure S2.** Confirming MIDAL with an alternative protease 11
**Figure S3.** Evaluating cleavage specificity for the Proline Endopeptidase 12
**Figure S4.** Confirming MIDAL with a photolabile linker 13
**Figure S5.** Crosslinking AGE intensities decrease after digestion 14
**Figure S6.** $^{1}$H NMR and COSY of MIDAL 15
**Figure S7.** $^{13}$C NMR and HSQC of MIDAL 16
**Figure S8.** Optimization of MIDAL formation 17
**Figure S9.** Evaluating MIDAL Stability 18

**General Materials.** Chemical reagents and solvents were of analytical grade, obtained from commercial suppliers without further purification unless otherwise specified. Methylglyoxal (MGO) (40% w/v in water) (M0252), Proline Specific Endopeptidase from Flavobacterium sp. (E1411) and TEV Protease (T4455) were purchased from MilliporeSigma. HEK-293T cells were purchased from ATCC. Rat α-GFP antibody (3H9) and GFP-Trap® Magnetic Agarose (GTMA) were purchased from Chromotek. α-rat secondary antibody (7077S) was purchased from Cell Signaling Technology. All Fmoc-protected amino acid monomers were purchased from ChemPrep Inc, or Advanced ChemTech Inc. Deuterium oxide for NMR analysis was purchased from Cambridge Isotope Laboratories, Inc. (DLM-4-10X0.7). Acetone-$D_6$ for $D_4$-MGO synthesis was purchased from Cambridge Isotope Laboratories, Inc. (DLM-9TC-10X0.5).

**Peptide Synthesis.** Peptides were prepared using standard Fmoc-based solid phase peptide synthesis on Fmoc-Ala-Wang resin (100-200 mesh, 0.64 mmol/g loading, CreoSalus Inc.) on a 100 μmol scale in a 10 mL polypropylene fritted syringe. Amino acids were added to the resin after initial Fmoc-deprotection (4 mL of 20% piperidine in DMF 2 x 15 min) and washes with DMF (4 mL, 4 x 1 min). Coupling was achieved by reacting 5 equiv. of the amino acid (relative to resin loading) in combination with 5 equiv. of *O*-(benzotriazol-1-yl)-*N,N,N',N'*-tetramethyluronium hexafluorophosphate (HBTU) and 10 equiv. of *N,N*-Diisopropylethylamine (DIEA) for 1 hour in a total of 3 mL of DMF. Amino acids following proline residues were double coupled to ensure high yield. Coupling was followed by subsequent DMF wash, fmoc-deprotection and another DMF wash before the next amino acid coupling. N-terminal acetylation was achieved by reacting 4 equiv. of acetic anhydride with 4 equiv. DIEA for 2 h in 3 mL of DMF. The sidechain protecting groups used were as follows: Arg(Pbf), Asn(Trt), Asp(tBu), Cys(Trt), Gln(Trt), Glu(tBu), His(Trt), Ser(Trt), Thr(tBu), and Tyr(tBu). Side-chain acid-deprotection and peptide cleavage was conducted with trifluoroacetic acid (TFA), triisopropylsilane (TIPS) and water (95:2.5:2.5) in 4 mL of acid solution. The resulting solution was then concentrated under air flow and re-dissolved in 2 mL of 50:50 water/acetonitrile mixture prior to purification.

**Peptide Purification.** Peptides were purified on a semi-preparative scale using an Agilent ZORBAX SB-C18 column (9.4 x 250 mm, 5 μm particle size) on an Agilent 1260 Infinity LC system with a water/acetonitrile mobile phase containing 0.1% TFA. 200 μL of crude peptide solution was injected onto the column and eluted using a standard purification gradient of 5-35% acetonitrile in water over 20 min with a flow rate of 3.5 mL/min. Absorbances were monitored at 215 and 280 nm and peaks were collected using an automated fraction collector. Collected fractions were characterized using a Bruker Microflex matrix-assisted laser desorption/ionization time-of-flight (MALDI-TOF) mass spectrometer. Fractions containing pure peptide of interest were combined and lyophilized. Peptides were reconstituted as 20 mM stocks in 50:50 DMF and water solutions.

**MALDI Mass Spectrometry.** Matrix-assisted laser desorption/ionization time-of-flight (MALDI-TOF) was used to assess peptides from HPLC purifications. Samples were co-crystallized using 0.2 μL of sample and 0.8 μL of matrix (α-cyano-4-hydroxycinnamic acid in 50% acetonitrile, 50% water with 0.1% trifluoroacetic acid) on ground steel plates.

**Liquid Chromatography-Mass Spectrometry.** Reversed phase liquid chromatography and mass spectrometry (LC-MS) analysis was carried out using an Agilent 1260 Infinity LC system coupled with Agilent 6530 Accurate Mass Q-TOF. Peptide reaction mixtures were injected onto an AdvanceBio Peptide 2.7 μm column (2.1 x 150 mm, Agilent) and eluted with a binary mobile phase of water with 0.1% formic acid (A) and acetonitrile with 0.1% formic acid (B). The elution method was as follows: isocratic 5% B from 0-1.75 min (0.400 mL/min); a gradient change to 15% B at 1.76 min followed by a change of 15% B to 40% B from 1.75-12.00 min (0.400 mL/min); column was then washed with 100% B and re-equilibrated to 5% B over the next 10 min with column heating at 55 °C. Mass spectrometry was accomplished with an electrospray ionization (ESI) source in the positive mode and spectra in the range of 300-3000 (*m/z*) were collected at a rate of 5 scans/sec from 1.75-20.00 min of the chromatography method. MS acquisition was achieved using the parameters: ESI capillary voltage, 4000 V; fragmentor, 150 V; gas temperature, 325 °C; gas rate, 12.5 L/min; nebulizer, 40 psig. When relevant, precursor ions were automatically selected for tandem mass spectrometry (collision induced dissociation) to generate MS/MS fragmentation spectra. Automated selection was based on both absolute (>200 counts) and relative (>0.01% of total counts) per cycle. Targeted MS/MS was performed on selected adducts, with their predicted *m/z* and retention time used to target them (Δ 0.2 min for rt and 4 *m/z* for isolation width). MS/MS spectra were acquired at a rate of 3 scans/sec using both a ramped collision energy model with a slope of 3.6 V and an offset of -4.8 V, a process that increases collision energy based on precursor ion drift time and a linear collision energy model that held the collision energy at 20 V for all precursor ions.

**LC-MS Analysis and Quantification.** Analysis of mass spectrometry data was carried out using Agilent MassHunter Qualitative Analysis software. MS data was quantified using the molecular feature extractor which reports cumulative ion counts (MS) as 'volumes' (size(*m/z*) x peak area) observed for all charge states associated with an ion. As peptides and their modified counterparts can ionize differently, this allows for more robust data than comparing individual charge states. For each peptide sample, compound lists were generated for each replicate of MGO treatment. Quantification was carried out by dividing the AGE adduct volume by the total volume of both modified and unmodified peptide. For digested samples, remaining adducts were divided by the total volume found in the corresponding intact run to find the %crosslink, as there were no unmodified full-length species left in the digested runs. This quantification allows for comparison of glycation across different peptides and glycating conditions. Additionally, absolute counts of peptides that were untreated showed similar levels of ionization. Retention times (RT) were used to identify discreate AGEs with degenerate masses. For tandem mass spectrometry, the collected scans were combined and reported for each identified precursor ion for each compound that triggered MS/MS acquisition. ProteinProspector (UCSF), an online proteomics tool, was used to generate a list of expected b and y ions for modified and unmodified peptides, which was used to assign the site of modification.

Percent glycation for ***intact*** runs was quantified according to the formula:

$$\%\,glycation = \frac{volume\ of\ AGE\ modified\ peptide}{total\ volume\ of\ both\ modified\ and\ unmodified\ peptide}$$

Percent glycation for crosslinks found in ***digested*** runs was quantified according to the formula:

$$\%\,crosslink = \frac{volume\ of\ AGE\ modified\ peptide\ remaining\ after\ digestion}{total\ volume\ of\ both\ modified\ and\ unmodified\ peptide\ from\ corresponding\ intact\ experiment}$$

**General Protocol for Glycation of Peptides.** Glycation reactions for peptides in solution were in general performed at a 20 µL scale in 200 µL Eppendorf tubes. MGO stocks (10 mM) were made fresh prior to each experiment and were prepared by diluting 15.4 µL of 40% w/v solution into 10 mL of ultrapure water. To perform *in vitro* glycation, 4 µL of 10X phosphate buffered saline (PBS) (1.37 M NaCl, 27 mM KCl, 100 mM $Na_2HPO_4$, 18 mM $KH_2PO_4$) at pH 7.4, 4 µL of 10 mM MGO stock in water, 11 µL of water and, finally, 1 µL of peptide stock in 50:50 DMF and water (20 mM). The final concentrations were 1 mM peptide, 2 mM MGO, 2X PBS with 2.5% DMF co-solvent. Reactions were pipetted up and down to ensure mixture of reactants. Tubes were capped and incubated in a 37 °C incubator, typically for 24 hours unless otherwise specified. After incubation, 3 µL aliquots were removed and the reaction was stopped by adding 1 µL of 500 mM Tris Buffer pH 7.3 (125 mM final concentration). Samples were then diluted in 297 µL of water unless otherwise noted and subjected to LC-MS analysis. For glycation reactions performed with glyoxal (GO), or deuterated MGO (D4-MGO) the same protocol was followed with GO final concentrations being 5 mM and D4-MGO at 2 mM.

**Crosslink Screening using Proline Protease.** To screen for crosslinking AGEs, an additional 3 µL aliquot was taken from glycation reactions. As before, 1 µL of 500 mM Tris Buffer pH 7.3 was added to stop the glycation reaction. An additional 1 µL of proline endopeptidase (0.05 units) was added and the resulting solution was incubated at 30°C for 15 min for complete digestion of peptide substrates. Samples were then diluted in 300 µL of water and subjected to LC-MS analysis.

**Crosslink Screening using Photocleavage.** To screen for crosslinking AGEs when the linker between the Arg residues was a photolabile linker, glycation reactions were performed as described above. After reaction, a 5 µL aliquot was quenched by adding 1.6 µL of 500 mM Tris Buffer pH 7.3 and diluted into 494 µL of water. Samples were then either subjected to LC-MS analysis (intact) or cleaved using UV radiation for 2 h with a power of 100 mwatts/cm$^2$ and 365 nm cutoff prior to LC-MS analysis.

**Methylglyoxal (MGO) Synthesis Protocol.** MGO was synthesized as described by Riley Oxidation.[5] To start, equal parts (10 mmol) deuterated acetone and selenium dioxide powder were added to water to a total volume of 5 mL in a round bottom flask with a magnetic stir bar. The reaction was heated in an oil bath at 100 °C under reflux for 4 hours. After reflux, distillation was performed by attaching the flask to a short path distillation head, and the mixture was distilled into one fraction at an internal temperature of 100 °C. The pale-yellow distillate was lyophilized overnight, revealing a viscous yellow MGO product (134 mg, 19%). Synthetic MGO was characterized by NMR, and a synthetic MGO stock was prepared by adding ultrapure water and checked for concentration using an aminoguanidine assay.[6,7] After a 5-hour incubation of aminoguanidine and MGO at 37 °C, absorbance was measured at 320 nm to generate a standard curve, the slope of which was compared to that of a calibration curve generated by serial dilutions of commercial MGO stock to equalize MGO concentrations between synthetic and commercial stocks.

**Preparation of MIDAL-modified Peptide 3.** For NMR studies, roughly 10 mg of $[M+144]_{XL}$-modified peptide **3** was prepared by incubating peptide **3** (250 µL of 20 mM stock) with MGO (2 mL of 20 mM stock) in 2x PBS (1 mL, pH 7.4 10x stock) and 1.75 mL of water, with a final concentration of 1 mM peptide **3**, 8 mM MGO, 2x PBS at pH 7.4 and 2.5% DMF in 5 mL total volume. This solution was incubated at 37 °C for 5 hours. The higher MGO concentration was used to promote crosslink formation. The resulting mixture of AGE-modified peptide was subsequently subjected to digestion with the proline endopeptidase 200 µL (10 units) for 30 min at 30 °C. This increased digestion time allowed for the use of less protease on large scale reactions. The $[M+144]_{XL}$ was purified by semi-preparative HPLC using a gradient of 5-25% acetonitrile in water over 40 min at 3.5 mL/min. Collected fractions were characterized by LCMS, pooled and lyophilized. This process was repeated until sufficient quantities of peptide **3** with $[M+144]_{XL}$ were obtained. Stocks were assessed for purity and protease resistance by LC-MS prior to analysis by NMR.

**NMR acquisition.**, $^{13}$C, COSY and HSQC spectra were acquired with a Bruker Advance III (500 MHz, 125 MHz) spectrometer. $^1$H chemical shifts are reported as units of parts per million (ppm) relative to $D_2O$(s 4.7 ppm). Data are reported as follows: chemical shift, multiplicity (s = singlet, d = doublet, t = triplet, q = quartet, m = multiplet), coupling constants (Hz), and integration. For NMR analysis, lyophilized $[M+144]_{XL}$ modified peptide **3** (10 mg) was dissolved in 550 µL of $D_2O$ and characterized by NMR. $^1$H NMR (500 MHz, D2O) δ 7.04 (dt, $J$ = 8.9, 2.4 Hz, 2H), 6.78 – 6.71 (m, 2H), 5.04 – 4.92 (m, 1H), 4.43 (dt, $J$ = 9.3, 4.8 Hz, 1H), 4.36 (s, 1H), 4.29 (dd, $J$ = 10.3, 4.9 Hz, 3H), 4.20 (s, 2H), 4.06 (d, $J$ = 17.3 Hz, 1H), 3.98 (d, $J$ = 16.9 Hz, 1H), 3.89 – 3.78 (m, 2H), 3.56 (td, $J$ = 13.1, 5.9 Hz, 2H), 3.32 – 3.26 (m, 3H), 3.15 (s, 1H), 2.91 (dd, $J$ = 13.9, 6.8 Hz, 1H), 2.86 (s, 2H), 2.19 (dt, $J$ = 12.4, 7.9 Hz, 1H), 1.96 (q, $J$ = 6.7 Hz, 1H), 1.95 – 1.87 (m, 1H), 1.89 (s, 3H), 1.79 (s, 1H), 1.67 (s, 2H), 1.58 – 1.40 (m, 18H), 1.40 – 1.27 (m, 4H), 0.88 – 0.73 (m, 12H).

**Mammalian cell culture.** HEK293-T cells (ATCC, CRL-3216) were cultured under standard conditions in Dulbecco's Modified Eagle Medium (DMEM) supplemented with 10% Fetal Bovine Serum (FBS) and 1% Penicillin-Streptomycin (Pen-Strep) at 37 °C at 5% CO2. Cells were passaged every 2 to 3 days for a maximum of 20-25 passages.

**Plasmids for Protein Expression.** Plasmids encoding green fluorescent protein (GFP) variants C-terminally fused to peptide 3 (-YLRGPGRLA, GFP-**3**), peptide 6 (-YLCGPGRLA, GFP-**6**), peptide **7** (-YLKGPGRLA, GFP-7) were designed and purchased from ThermoFisher GeneArt, in a pcDNA_3.1(+) vector, which was selected for its propensity for uninduced, high expression in mammalian cell lines. The C-terminal peptide sequences were connected to GFP through a linker sequence containing a tobacco etch virus (TEV) protease cleavage site. The expression sequences used to encode the GFP-fusion proteins are shown below. All sequences contain the sequence for green-fluorescent protein (bp 1-714) linked via the TEV cleavage site (bp 715-735) to the peptide of interest (bp 736-762). The peptide sequence is the only variable region between plasmid sequences.

GFP-**3**:

ATGAGCAAGGGCGAAGAACTGTTCACCGGCGTGGTGCCCATTCTGGTGGAACTGGACGGGGATGT
GAACGGCCACAAGTTTAGCGTTAGCGGCGAAGGCGAAGGGGATGCCACATACGGAAAGCTGACC
CTGAAGTTCATCTGCACCACCGGCAAGCTGCCTGTGCCTTGGCCTACACTGGTCACCACCTTTACCT
ACGGCGTGCAGTGCTTCAGCAGATACCCCGACCATATGAAGCGGCACGACTTCTTCAAGAGCGCC
ATGCCTGAGGGCTACGTGCAAGAGCGGACCATCTTCTTTAAGGACGACGGCAACTACAAGACCAG
GGCCGAAGTGAAGTTCGAGGGCGACACCCTGGTCAACCGGATCGAGCTGAAGGGCATCGATTTCA
AAGAGGACGGCAACATCCTGGGCCACAAGCTTGAGTACAACTACAACAGCCACAACGTGTACATC
ATGGCCGACAAGCAGAAAAACGGCATCAAAGTGAACTTCAAGATCCGGCACAACATCGAGGACG
GCTCTGTGCAGCTGGCCGATCACTACCAGCAGAACACACCCATCGGAGATGGCCCTGTGCTGCTGC
CCGATAACCACTACCTGAGCACACAGAGCGCCCTGAGCAAGGACCCCAACGAGAAGAGGGATCA
CATGGTGCTGCTGGAATTTGTGACCGCCGCTGGCATCACCCACGGCATGGATGAGCTGTACAAAG
AGAACCTGTACTTCCAGAGCTACCTGAGAGGCCCTGGCAGACTGGCTTAA

GFP-**6**:

ATGAGCAAGGGCGAAGAACTGTTCACCGGCGTGGTGCCCATTCTGGTGGAACTGGACGGGGATGT
GAACGGCCACAAGTTTAGCGTTAGCGGCGAAGGCGAAGGGGATGCCACATACGGAAAGCTGACC
CTGAAGTTCATCTGCACCACCGGCAAGCTGCCTGTGCCTTGGCCTACACTGGTCACCACCTTTACCT
ACGGCGTGCAGTGCTTCAGCAGATACCCCGACCATATGAAGCGGCACGACTTCTTCAAGAGCGCC
ATGCCTGAGGGCTACGTGCAAGAGCGGACCATCTTCTTTAAGGACGACGGCAACTACAAGACCAG
GGCCGAAGTGAAGTTCGAGGGCGACACCCTGGTCAACCGGATCGAGCTGAAGGGCATCGATTTCA

AAGAGGACGGCAACATCCTGGGCCACAAGCTTGAGTACAACTACAACAGCCACAACGTGTACATC
ATGGCCGACAAGCAGAAAAACGGCATCAAAGTGAACTTCAAGATCCGGCACAACATCGAGGACG
GCTCTGTGCAGCTGGCCGATCACTACCAGCAGAACACACCCATCGGAGATGGCCCTGTGCTGCTGC
CCGATAACCACTACCTGAGCACACAGAGCGCCCTGAGCAAGGACCCCAACGAGAAGAGGGATCA
CATGGTGCTGCTGGAATTTGTGACCGCCGCTGGCATCACCCACGGCATGGATGAGCTGTACAAAG
AGAACCTGTACTTCCAGAGCTACCTGTGTGGCCCTGGCAGACTGGCTTAA

GFP-**7**:

ATGAGCAAGGGCGAAGAACTGTTCACCGGCGTGGTGCCCATTCTGGTGGAACTGGACGGGGATGT
GAACGGCCACAAGTTTAGCGTTAGCGGCGAAGGCGAAGGGGATGCCACATACGGAAAGCTGACC
CTGAAGTTCATCTGCACCACCGGCAAGCTGCCTGTGCCTTGGCCTACACTGGTCACCACCTTTACCT
ACGGCGTGCAGTGCTTCAGCAGATACCCCGACCATATGAAGCGGCACGACTTCTTCAAGAGCGCC
ATGCCTGAGGGCTACGTGCAAGAGCGGACCATCTTCTTTAAGGACGACGGCAACTACAAGACCAG
GGCCGAAGTGAAGTTCGAGGGCGACACCCTGGTCAACCGGATCGAGCTGAAGGGCATCGATTTCA
AAGAGGACGGCAACATCCTGGGCCACAAGCTTGAGTACAACTACAACAGCCACAACGTGTACATC
ATGGCCGACAAGCAGAAAAACGGCATCAAAGTGAACTTCAAGATCCGGCACAACATCGAGGACG
GCTCTGTGCAGCTGGCCGATCACTACCAGCAGAACACACCCATCGGAGATGGCCCTGTGCTGCTGC
CCGATAACCACTACCTGAGCACACAGAGCGCCCTGAGCAAGGACCCCAACGAGAAGAGGGATCA
CATGGTGCTGCTGGAATTTGTGACCGCCGCTGGCATCACCCACGGCATGGATGAGCTGTACAAAG
AGAACCTGTACTTCCAGAGCTACCTGAAAGGCCCTGGCAGACTGGCTTAA

**Cellular MGO Assay (Treatment A).** For experiments to evaluate glycation of each GFP variant, roughly 2.6 million HEK-293T cells were seeded in a 10 cm sterile tissue culture dish and grown to 50-60% confluency (~24 hours). They were then transfected with 10 μg of the desired plasmid using TransIT-LT1 Transfection Reagent (MirusBio, 2 μL/μg plasmid). After transfection, cells were cultured for an additional 24 h (to >90% confluency) before MGO treatments and subsequent harvesting. In general, for "treatment A" cells were treated with MGO diluted into DMEM supplemented with 10% FBS and 1% Pen-Strep, in stocks that were prepared the same day as MGO treatment. MGO stock solutions (100 mM) were prepared in sterile water by the addition of 154 μL of commercially available MGO (40% w/v) to a total volume of 10 mL. The 100 mM MGO stock was further diluted into DMEM until a desired concentration (10 mM final concentration) was reached. Adherent cells on 10 cm sterile culture dishes were treated with or without 10 mL of this MGO-supplemented DMEM and incubated at 37 °C at 5% $CO_2$ for 2 h. After MGO treatment, cells were harvested with TrypLE Express (Gibco). Harvested cells were transferred to a 15 mL conical tube, and pelleted at 200 x g. The resulting pellet was washed with 5 mL of 20 mM PBS, pH 7.3 and then lysed on ice in 400 μL Tris-Cl buffer (50 mM Tris, 150 mM NaCl, 1 mM EDTA, 1mM NaF, 1% Triton X-100) at pH 7.5 with a Pierce Protease and Phosphatase Inhibitor tablet (1 tablet/10 mL buffer). Lysates were clarified by centrifugation at 17000 x g for 10 minutes, and total protein quantified by BCA Protein Assay (Pierce) on a Tecan Spark 10M plate reader and analyzed by western blot (10 μg/sample) or used for immunoprecipitation.

**Immunoprecipitation Protocol.** Cellular lysates (200-250 μg protein) were diluted to a total of 500 μL in Eppendorf tubes containing 10 mM Tris/Cl pH 7.5, 150 mM NaCl, 0.5 mM EDTA. This diluted lysate was added to 25 μL of equilibrated GFP-Trap Magnetic Agarose beads and allowed to incubate for 2 h at 4 °C with end over end rotation. Following incubation, the beads were separated with a magnet until the supernatant was clear. The beads were washed 3 times with 500 μL of the same dilution buffer and transferred to a new tube. For on bead digestion with TEV proteolysis, the beads were then resuspended in 21 μL of dilution buffer and 4 μL (40 units) of TEV protease (MilliporeSigma) were added and allowed to incubate at room temperature for 2 hours with end over end rotation. Following incubation, the supernatant was collected and subjected to LC-MS analysis or further proline endopeptidase cleavage and then LC-MS analysis.

**Extracellular MGO Assay (Treatment B).** As a control, MGO "treatment B" was conducted after immunoprecipitation of cellular lysates, but prior to TEV cleavage. Untreated lysates were pulled down using the above protocols. Once the beads were washed, incubated with lysate and washed again, they were subsequently washed 3 times with 2x PBS. They were then resuspended in 50 μL of 2x PBS with 2 mM MGO. Samples were incubated on a shaker at 37 °C for 5 h before they were washed 3 more times with dilution buffer prior to proceeding with the protocol (described above) to cleave and analyze the samples.

**Western Blotting.** Lysates with a concentration of 10 μg total protein were diluted with 6X SDS loading buffer and boiled for 5 min. SDS-PAGE analysis was performed using pre-cast protein gels (8-16%, mini-PROTEAN TGX) in standard Tris/glycine/SDS running buffer at 200 mV for 40 min to resolve protein bands. Proteins were transferred to PVDF membrane using an iBlot 2 (Invitrogen). Membranes were then blocked in buffer containing 20 mM Tris, 150 mM NaCl, 0.1% Tween (1x TBST) with 5% w/v BSA. After blocking, primary antibody was incubated overnight at 4 °C. Membranes were subsequently washed for 3 x 5 minutes with the same TBST with 5% BSA buffer. Secondary HRP-conjugated antibodies were then incubated with the membranes in TBST with 5% BSA for an hour at room temperature. Membranes were then again washed 3 times, developed using Clarity Western ECL Substrate (Bio-Rad) and imaged on a Bio-Rad ChemiDoc XRS+.

| Peptide | Mass (Da) | ppm error | RT (min) |
|---|---|---|---|
| **1** | 986.5658 | 0.4 | 8.386 ± 0.027 |
| **2** | 986.5653 | 0.9 | 8.465 ± 0.043 |
| **3** | 1043.587 | 0.6 | 7.846 ± 0.018 |
| **4** | 1100.6048 | 3.9 | 8.217 ± 0.020 |
| **5** | 1214.6502 | 1.5 | 10.560 ± 0.024 |
| **6** | 990.5003 | 4.6 | 9.767 ± 0.062 |
| **7** | 1015.5794 | 2.1 | 7.751 ± 0.027 |
| **S1** | 2079.0286 | 0.03 | 12.920 ± 0.027 |
| **S2** | 874.3624 | 5.1 | 10.240 ± 0.035 |
| **S3** | 916.4387 | 1.8 | 9.511 ± 0.058 |
| **S4** | 1112.5797 | 5.6 | 7.238 ± 0.006 |
| **GFP-3, after TEV proteolysis** | 1088.6078 | 1.2 | 6.566 ± 0.016 |
| **GFP-6, after TEV proteolysis** | 1035.5155 | 1.6 | 8.014 ± 0.034 |
| **GFP-7, after TEV proteolysis** | 1060.6088 | 5.5 | 6.492 ± 0.018 |

**Table S1. Peptide masses and retention times.** Listed are the peptides that were used in this study. Observed masses (Da) with error from expected (ppm) along with retention times (min) including standard deviation are shown.

| Adducts | Mass (Da) | ppm error | RT intact (min) | RT digest(min) |
|---|---|---|---|---|
| **3**$^{MGH-1}$ | 1097.5967 | 1.4 | 8.112 ± 0.014 | n/a |
| **3**$^{MGH-DH}$ | 1115.6092 | 0.3 | 7.860 ± 0.013 | n/a |
| **3**$^{CEA}$ | 1115.6091 | 0.3 | 8.249 ± 0.015 | n/a |
| **3**$^{M+108}$ | 1151.6046 | 3.7 | 8.384 ± 0.013 | n/a |
| **3**$^{M+216}$ | 1169.6148 | 3.9 | 8.117 ± 0.006 | n/a |
| **3**$^{M+216}$ | 1169.6160 | 2.9 | 8.312 ± 0.014 | n/a |
| **3**$^{THP}$ | 1187.6326 | 2.2 | 7.913 ± 0.104 | n/a |
| **3**$^{M+144}$ | 1187.6338 | 3.2 | 8.231 ± 0.014 | n/a |
| **3**$^{M+144}$ | 1187.6322 | 1.9 | 8.412 ± 0.009 | n/a |
| **3**$^{MIDAL}$ | 1187.6321 | 1.8 | 7.855 ± 0.012 | 7.858 ± 0.042 |
| **3**$^{M+198}$ | 1241.6393 | 1 | 8.260 ± 0.015 | n/a |
| **3**$^{M+216}$ | 1259.6488 | 1.8 | 8.655 ± 0.009 | n/a |
| **3**$^{M+216}$ | 1259.6489 | 1.7 | 8.180 ± 0.009 | 8.188 ± 0.044 |
| **3**$^{M+216}$ | 1259.6479 | 2.5 | 8.381 ± 0.004 | 8.38 ± 0.047 |
| **3**$^{M+288}$ | 1331.6658 | 4.8 | 8.355 ± 0.035 | n/a |
| **GFP-3**$^{MIDAL}$ | 1232.6555 | 3.3 | 6.616 ± 0.009 | 6.611 ± 0.012 |
| **6**$^{MICA}$ | 1026.4940 | 1.7 | 10.260 ± 0.009 | 10.29 ± 0.058 |
| **GFP-6**$^{MICA}$ | 1071.5137 | 3.2 | 8.395 ± 0.023 | 8.372 ± 0.041 |
| **7**$^{MODIC}$ | 1051.5766 | 4.7 | 9.907 ± 0.025 | n/a |

**Table S2. AGE adducts observed.** Complete list of AGEs observed after MGO glycation reactions with peptide **3.** For peptide **6,** or peptide **7**, or the peptides released from GFP-3 or GFP-6 after TEV proteolysis, only xl-AGEs are provided here. For each adduct, the observed mass (Da) and error (ppm) from expected mass are provided, along with the retention times ± standard deviations found in intact and, when relevant, digested samples.

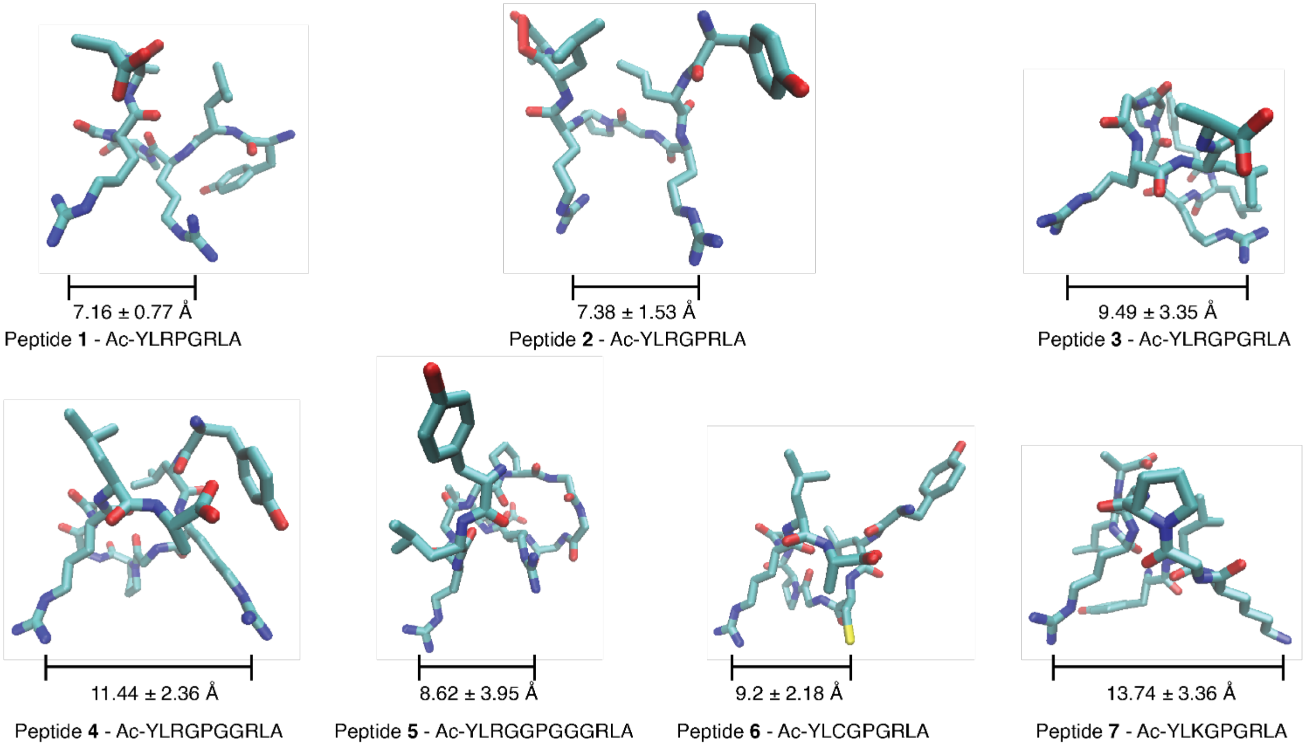


**Figure S1. Alpha fold predictions of peptides 1—7.** Representative conformations for peptides **1**—**7** predicted by AlphaFold. The guanidino groups for peptides **1**—**5** are facing each other in all predictions. The average (mean) distance between reactive residues (guanidino carbon for arginine, ε-amino group for lysine and thiol of cysteine) as well as the standard deviation calculated from the five most common conformations found using AlphaFold. Of note is that peptide **7** has a much larger distance between glycation sites than the other peptides, which may contribute to the minimal amounts of MODIC found during glycation reactions.

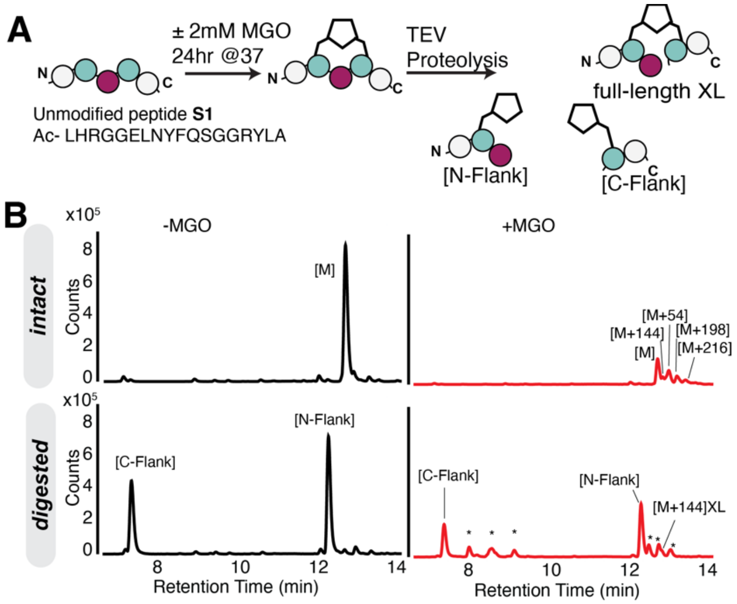


**Figure S2. Confirming MIDAL with an alternative protease. A)** Scheme depicting a control experiment using an alternative protease. Peptide **S1** contained a tobacco etch virus (TEV) protease recognition site to ensure that the MIDAL was not an artifact of the proline endopeptidase. **B)** Peptide **S1** was incubated under standard glycating conditions, samples were split, half analyzed by LC-MS (intact) and the remaining half subjected to 1 µL (10 units) of TEV protease and incubated at 4 °C overnight (18-24 h). Following digestion, a full-length [M+144] $_{XL}$ mass adduct was observed, along with the expected unmodified N- and C- flank peptides and their mono-AGEs (*).

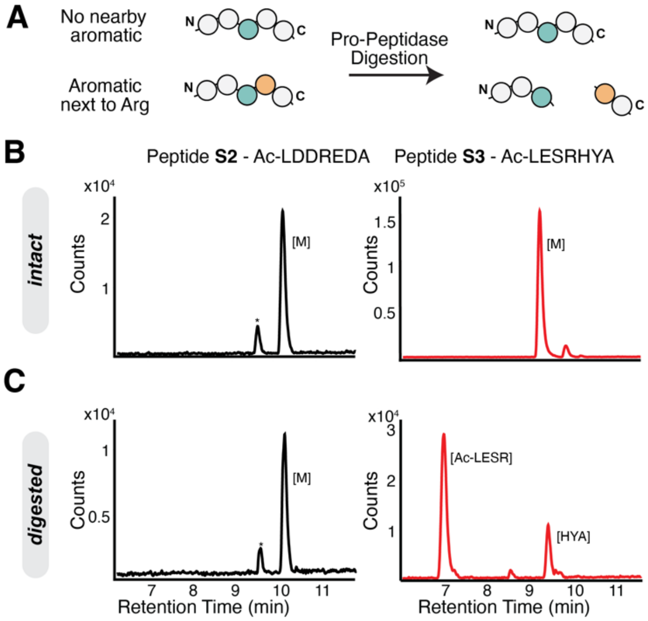


**Figure S3. Evaluating cleavage specificity for the Proline Endopeptidase**. Our platform for discovering xl-AGEs relies on a proline specific endopeptidase from Flavobacterium sp. (E1411), which we found to exhibit off-target cleavage when an arginine residue was followed by an aromatic residue (in this case histidine). **A)** We confirmed this behavior by incubating 1 mM control peptides (peptide **S2**: Ac-LDDREDA and peptide **S3**: Ac-LESRHYA) with the 0.05 units of the Pro peptidase for 15 min at 30 °C. **B)** Base peak chromatograms (BPC) for peptide **S1** (*left*) or **S2** (*right*) shown before and after treatment with the proline endopeptidase cleavage (intact and digested, respectively). For peptide **S1**, without an aromatic residue next to the Arg, only full-length peptide is observed, as would be expected for a peptide without any Pro. However, when an aromatic residue is introduced such as in peptide **S2** (right), complete cleavage of the peptide is observed after digestion with the proline endopeptidase. This experiment informed the design of the peptides we used in this study.

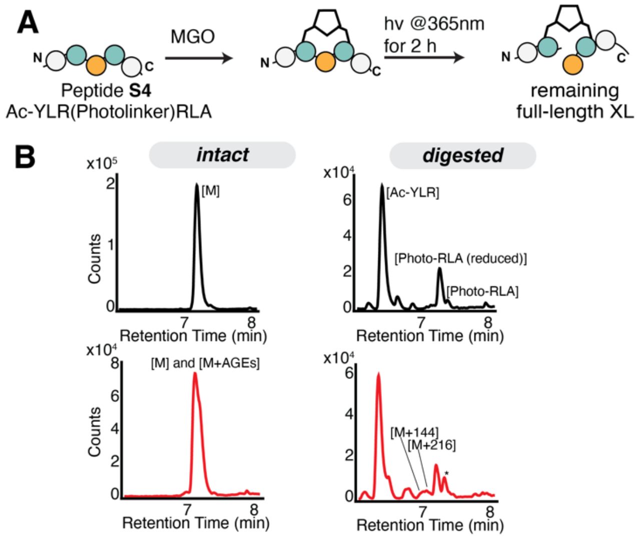


**Figure S4. Confirming MIDAL with a photolabile linker. A)** As an alternative to enzymatic cleavage, we prepared peptides containing a photo-cleavable ANP linker that could also reveal xl-AGE formation. Peptide **S4** was incubated under standard glycating conditions, samples were split, half analyzed by LC-MS (intact) and the remaining half was subjected to UV irradiation to cleave the peptide prior to LC-MS analysis (digested). **B)** Just as we observed with the enzymatic cleavage, the only full-length (or above) masses observed on the LC-MS after digestion were an $[M+144]_{XL}$ (MIDAL) and $[M+216]_{XL}$ adduct.

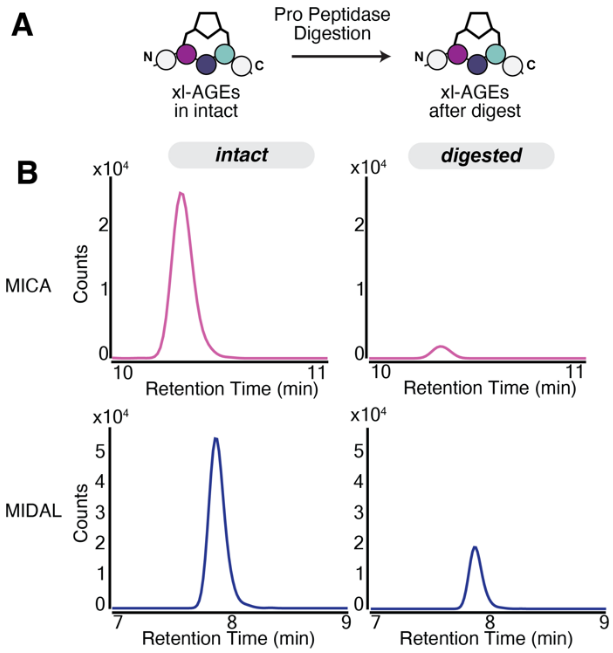


**Figure S5. Crosslinking AGE intensities decrease after digestion. A)** After digestion of peptide glycation reactions, a loss of counts for crosslinking adducts was observed. **B)** Representative extracted ion chromatograms for MICA (*left*) MIDAL (*right*) before and after treatment with the proline endopeptidase cleavage (intact and digested, respectively), with identical axes for intact and digested samples. Integration of peak volumes revealed more than a 15-fold decrease in peak volume for MICA (intact = 198037; digested = 12574). Similarly, MIDAL exhibited a 2-fold decrease in peak volume (intact = 304389; digested = 131375 post digestion. We suspect that the loss of counts can be attributed largely to a decrease in crosslink stability post-digestion. For this reason, quantification of xl-AGE levels was performed on intact samples, unless otherwise specified.

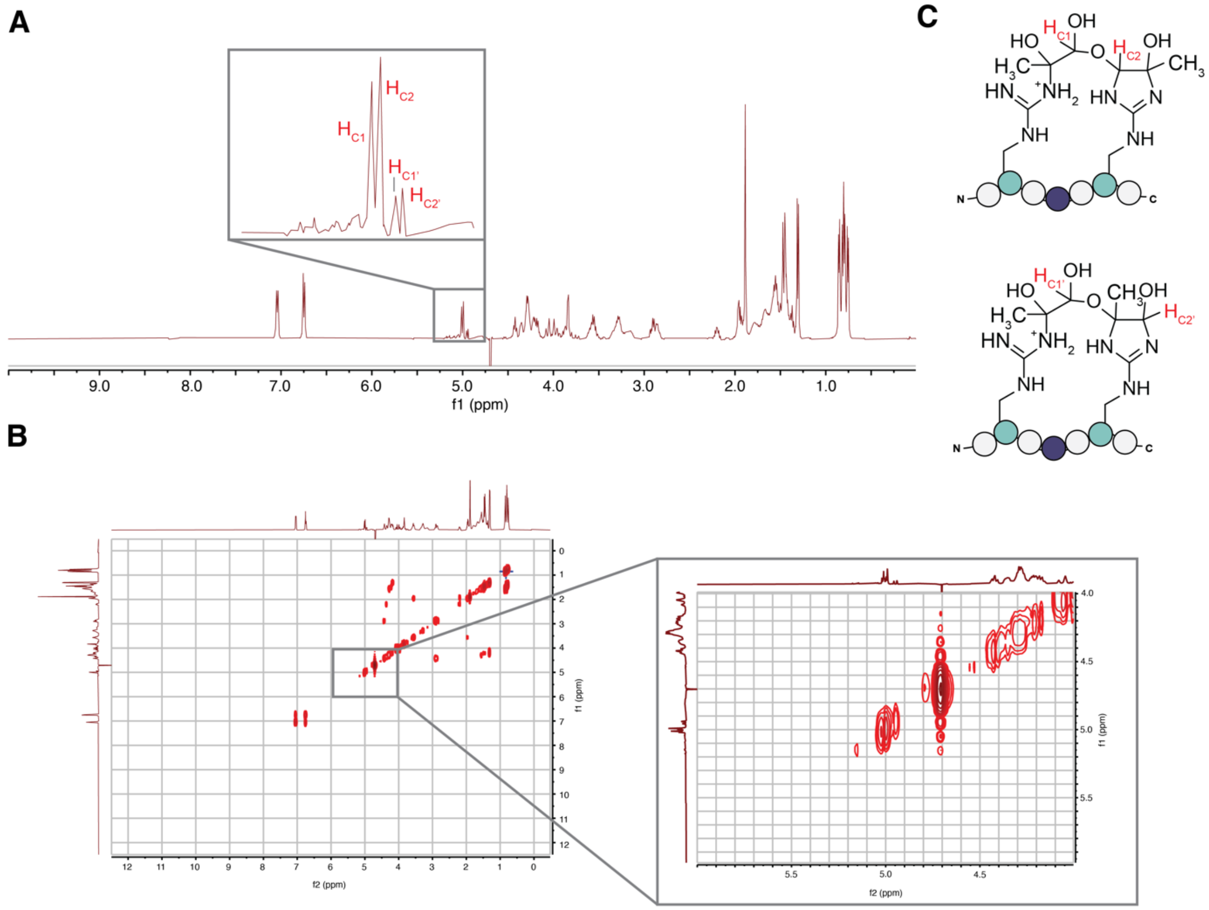


**Figure S6. $^1$H NMR and COSY of MIDAL.** Sufficient quantities of purified peptide **3$^{\mathbf{MIDAL}}$** were prepared to allow for structural studies by NMR. **A)** Using an HOD suppression experiment, a clear view of chemical shifts in the 4.5-5.5 ppm range was acquired. The $^1$H spectrum is shown, with key resonances highlighted 5.04 – 4.92 (m, 1H). These peaks show a multiplet, which would be expected for the predicted crosslinking structure. The ratio of MIDAL-1 to MIDAL-2 was calculated from the integration of peak $H_{C2}/H_{C2'}$ and found to be in a 3:1 ratio. Due to the spectra being taken in $D_2O$, chemical shifts for hydroxyl protons are not observed. **B)** Using COSY, a lack of cross peaks with other protons in the molecule is consistent with the proposed crosslink structure. **C)** These data allowed us to assign the $[144]_{XL}$ as a mix of two isomers (MIDAL-1, major 75%) and (MIDAL-2, minor 25%). Chemical shifts corresponding to the protons highlighted in red are labeled in panel A..

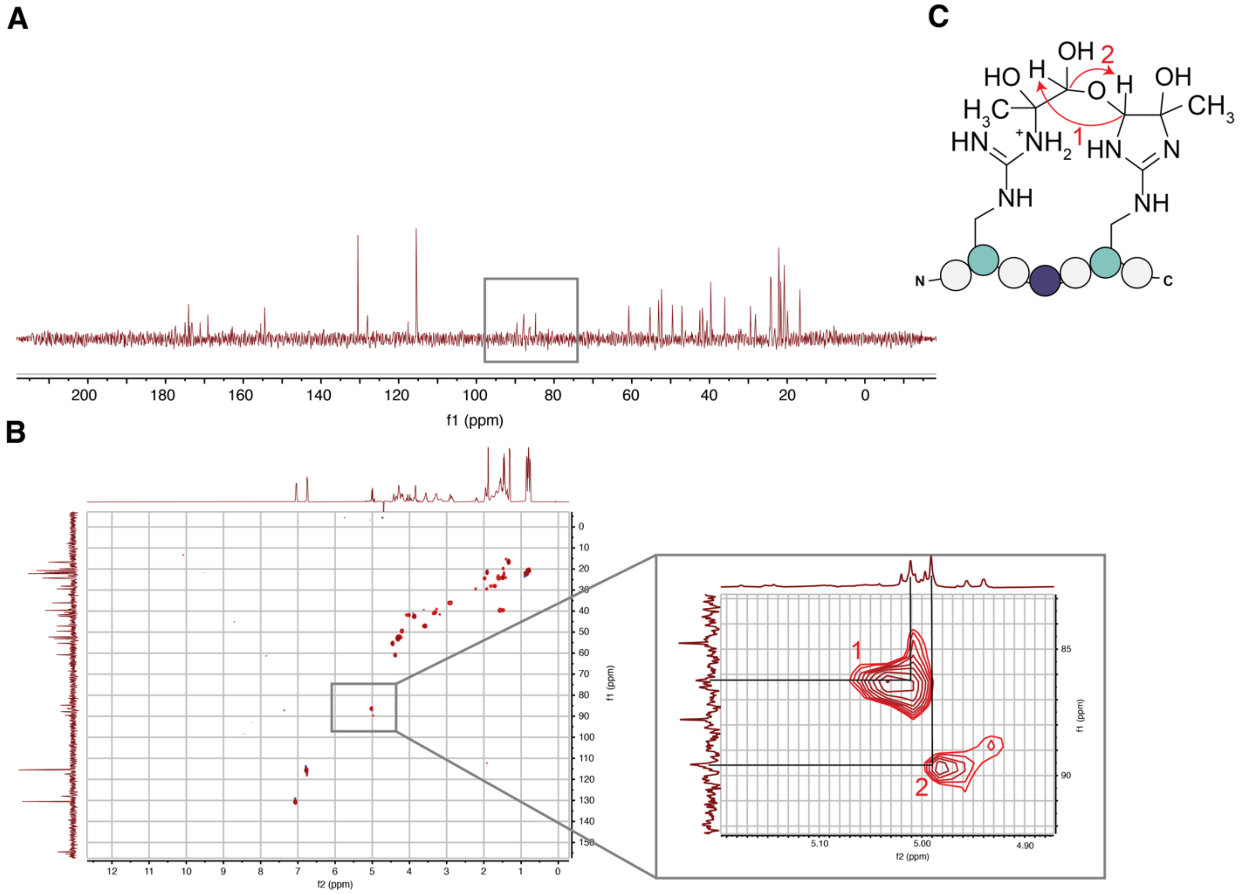


**Figure S7. $^{13}$C NMR and HSQC of MIDAL.** Sufficient quantities of purified peptide **3$^{MIDAL}$** were prepared to allow for structural studies by NMR. **A)** The $^{13}$C spectra shows clear chemical shifts between 80-90 ppm that were absent for the unmodified peptide. Peaks in this range are consistent with shifts expected for carbons participating in the hemiacetal group of the crosslink. **B)** HSQC acquired shows cross peaks that correspond to the major crosslinking product. **C)** MIDAL structure labeled with corresponding expected HSQC cross peak interactions.

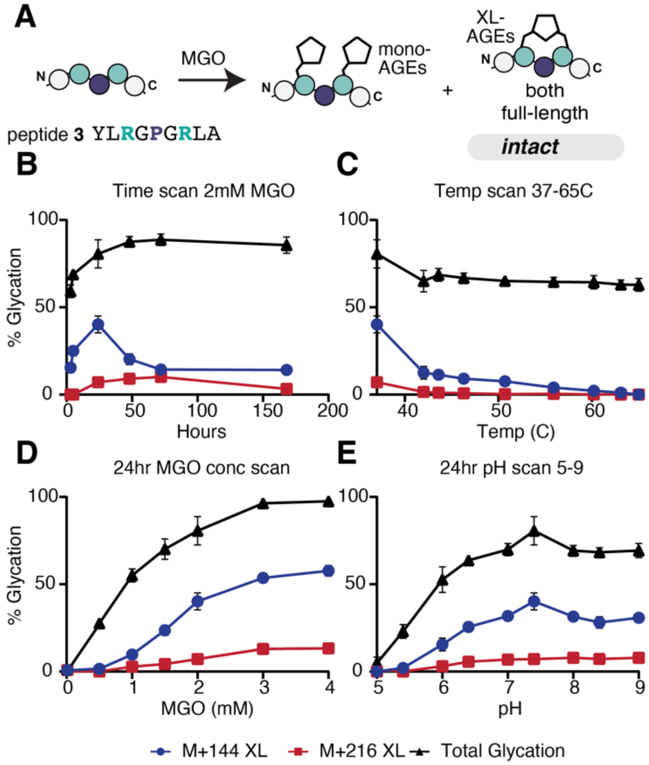


**Figure S8. Optimization of MIDAL formation. A)** To identify conditions that produce the greatest levels of MIDAL, peptide **3** (1 mM) was subjected to a variety of conditions and intact data was then analyzed by LCMS to track xl-AGE formation. **B)** AGE distributions shown for peptide **3** (1 mM) treated with 2 mM MGO for up to 1 week at 37 °C. n=3 for 3, 4, 48, 72 and 168 h and n=9 for the 24 h timepoint. These data show that MIDAL levels peak at 24 h. **C)** AGE distributions shown for peptide **3** (1 mM) treated with 2 mM MGO for 24 h at varying temp from 37 °C-65 °C. n=9 for 37 °C and n=3 for all other temperatures (42, 43.6, 46.3, 50.7, 55.9, 60.2, 63.1, 65 °C). These data show that MIDAL levels are greatest at physiological temperatures. **D)** AGE distributions shown for peptide **3** (1 mM) treated with a MGO concentrations ranging from 0-4 mM for 24 h at 37 °C. n=9 for 2 mM MGO and n=3 for all other MGO concentrations (0.5, 1, 1.5, 3 and 4 mM). These data show that MIDAL levels increase with overall glycation levels, but that a sharp increase is observed around 2 mM, which aligns with the expected stoichiometry for its formation. **E)** AGE distributions shown for peptide **3** (1 mM) treated with 2 mM MGO for 24 h at 37 °C in 2x phosphate buffered saline (PBS) at pH 5-9. n=9 for pH 7.4 and n=3 for all other pH. These data show that MIDAL formation is greatest at physiological pH. All plotted data values are shown as the mean ± standard deviation for each adduct. Legend: dark blue $[M+144]_{XL}$, dark red $[M+216]_{XL}$, black total glycation.

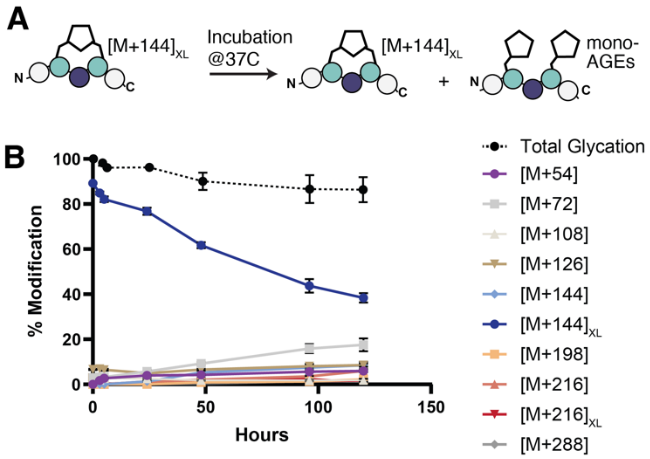


**Figure S9. Evaluating MIDAL Stability. A)** To assess MIDAL stability, purified peptide **3**[**MIDAL**] was incubated in 2X PBS at pH 7.4 for up to 5 days. The resulting AGE distributions were analyzed by LC-MS and quantified for each of the major adducts found (n=2). Legend: purple [M+54], light gray [M+72], light brown [M+108], brown [M+126], light blue [M+144], dark blue $[M+144]_{XL}$ (MIDAL), orange [M+198], light red [M+216], dark red $[M+216]_{XL}$, dark gray [M+288]. In general, over time MIDAL degraded fairly linearly, with a half-life of 80.4 ± 6.7 hours. As MIDAL levels decreased, we observed an increase in [M+72] levels, but all other AGEs remained constant. Levels of unmodified peptide **3** increased over the first 48 hours of incubation, after which time it leveled out, plateauing at 8.6 ± 3.7%. This indicates that MIDAL likely undergoes further rearrangement that breaks the hemiacetal, either leaving an [M+72] adduct on a single arginine or reverting to starting material. This data shows that MIDAL has an intermediate level of stability, suggesting it could be a reversible crosslink.
